## Supplemental Figures for "N-Terminus of *Drosophila Melanogaster* MSL1 Is Critical for Dosage Compensation"

**FIGURE SUPPLEMENT**

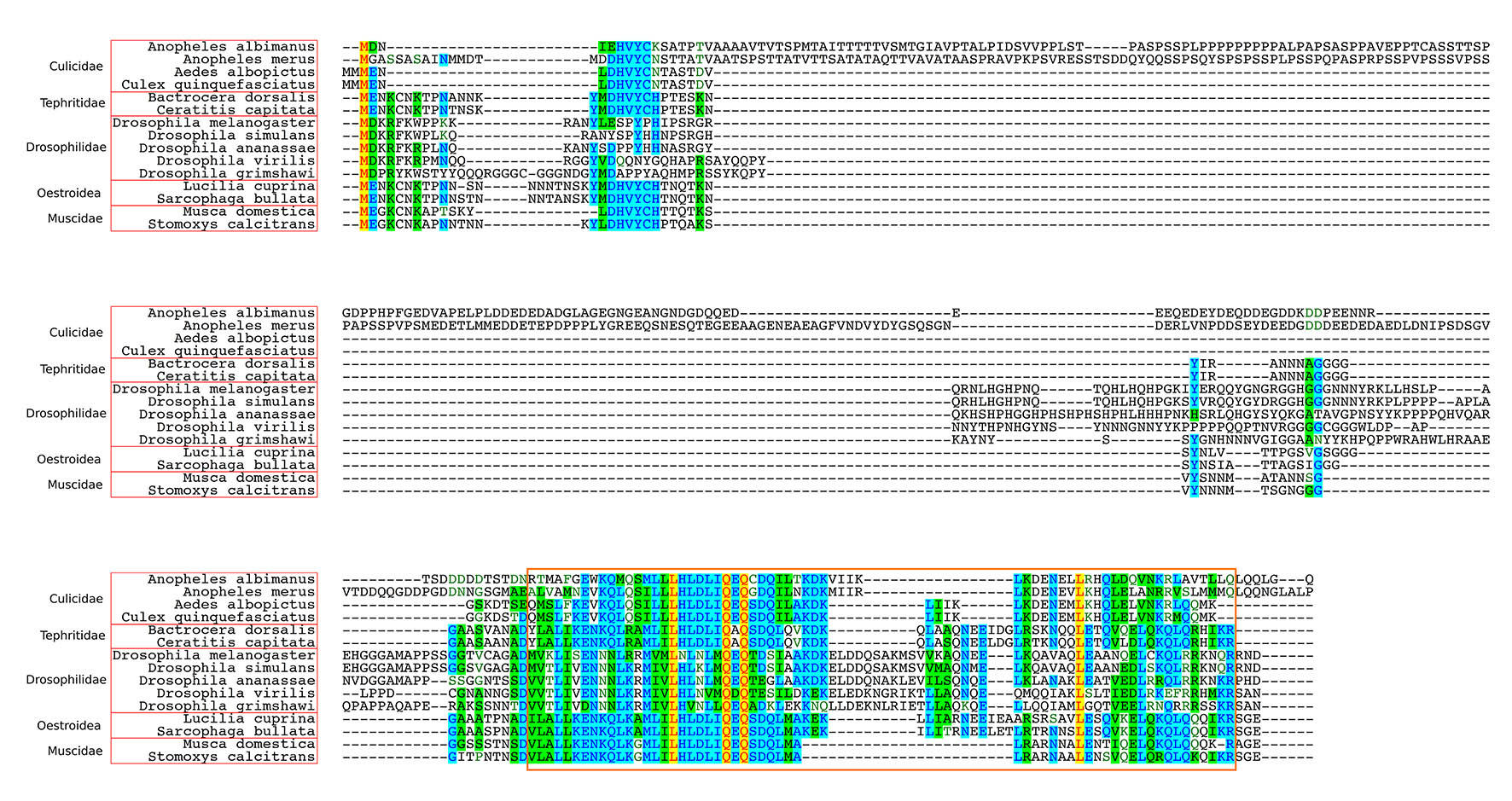

**Figure 1 – figure supplement 1.** Alignment of the N-terminal region of MSL1 among Diptera. Orange box – Coiled coil domain.

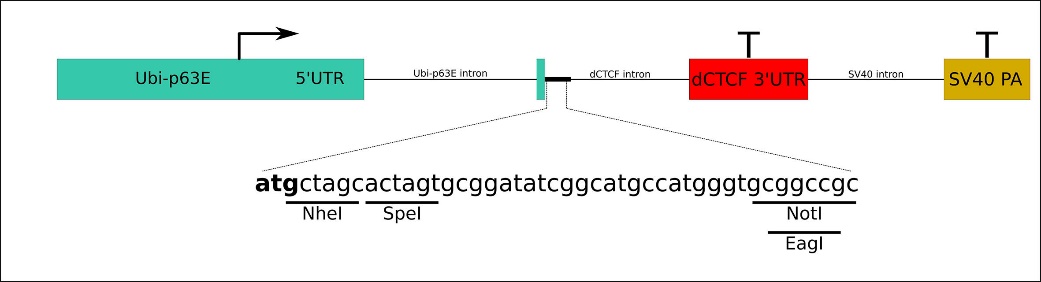

**Figure 1 – figure supplement 2.** Scheme of transgenic construct used to express MSL1 variants. Green boxes – promoter and 5’UTR of *Ubi-p63E* gene; red box – 3’UTR with polyadenylation signal from *dCTCF* gene; yellow box – polyadenylation signal from SV40 virus. Lines correspond to introns. Letters show the polylinker region used for cloning of MSL1 cDNAs.

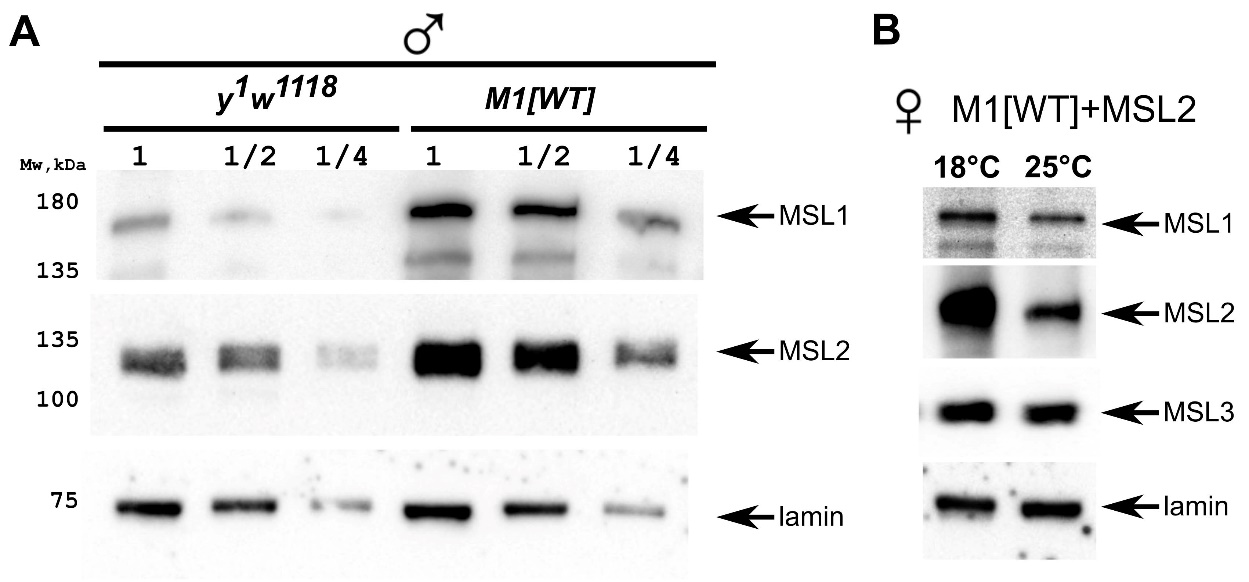

**Figure 1 – figure supplement 3. (A)** Comparing of MSL1 expression in *y^1^w^1118^* and M1[wt] (*msl-1*^−^*; Ubi:msl-1^WT^/ Ubi:msl-1^WT^*) males. Male samples were titrated as two-fold dilutions (‘1’, ‘1/2’, ‘1/4’). **(B)** Comparing of MSL1, MSL2, MSL3 expression in M1[wt]+MSL2 females (*msl-1*^−^*; Ubi:msl-2^WT^-FLAG* */ Ubi:msl-1^WT^*) growing at 18°C and 25°C. Immunoblot analysis performed using anti-MSL1, anti-MSL2, anti-MSL3, and anti-lamin Dm0 (internal control) antibodies.

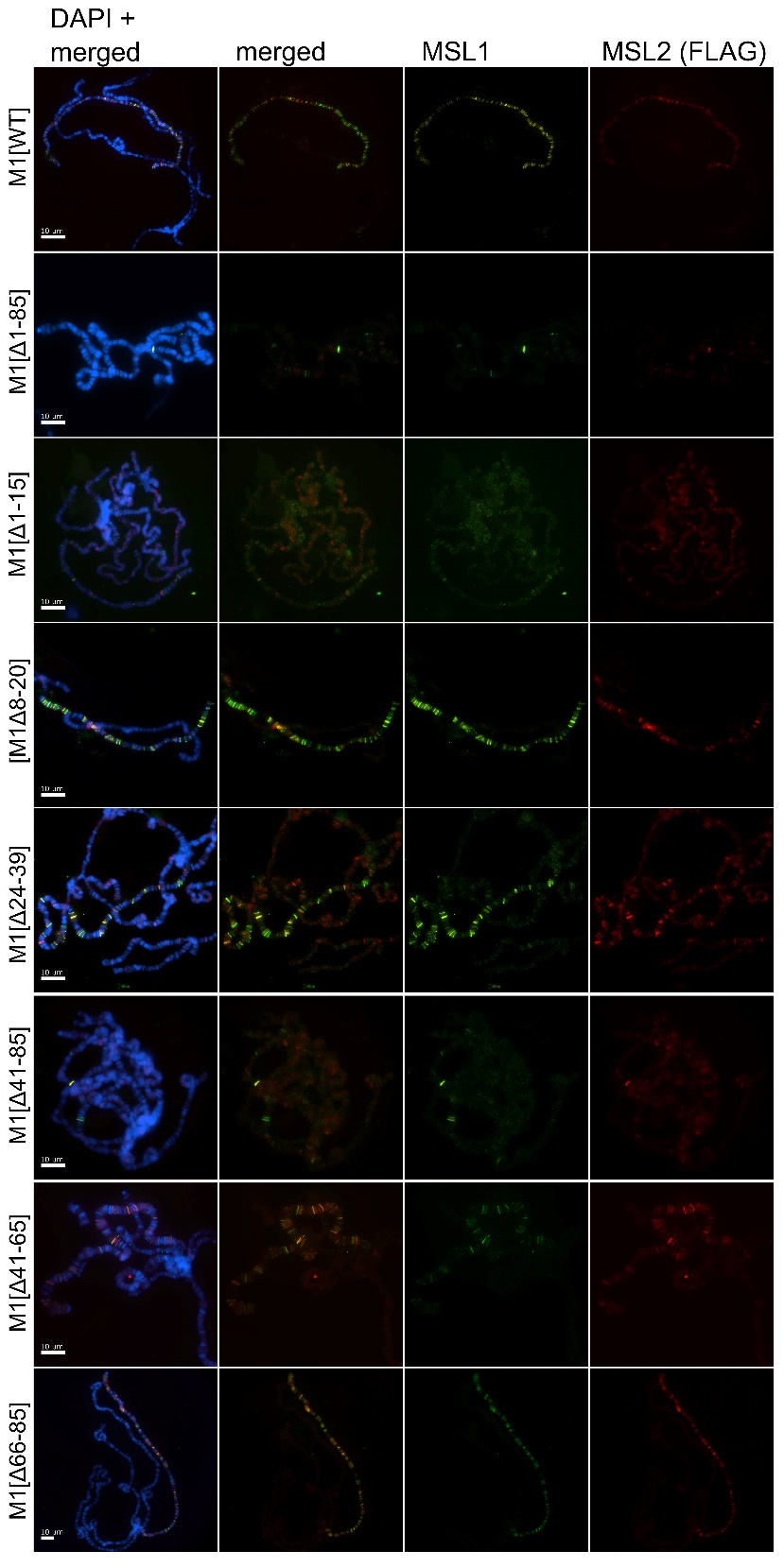

**Figure 2 – figure supplement 1. Testing the role of MSL1 mutants in recruiting the MSL complex on the X chromosome in a female model system.** Distribution of the MSL complex on the polytene chromosomes from 3rd-day female larvae expressing both MSL1 variants and the MSL2-FLAG protein. Panels show the immunostaining of proteins using rabbit anti-MSL1 antibody (green) and mouse anti-FLAG antibody (red). DNA was stained with DAPI (blue). M1[WT] (♀ *msl-1*^−^*; Ubi:msl-2^WT^-FLAG/ Ubi:msl-1^WT^*); M1[Δ1-85] (♀ *msl-1*^−^*; Ubi:msl-2^WT^-FLAG/ Ubi:msl-1^Δ1-85^*); M1[Δ1-15] (♀ *msl-1*^−^*; Ubi:msl-2^WT^-FLAG/ Ubi:msl-1^Δ1-15^*); M1[Δ8-20] (♀ *msl-1*^−^*; Ubi:msl-2^WT^-FLAG/ Ubi:msl-1^Δ8-20^*); M1[Δ24-39] (♀ *msl-1*^−^*; Ubi:msl-2^WT^-FLAG/ Ubi:msl-1^Δ24-39^*); M1[Δ41-85] (♀ *msl-1*^−^*; Ubi:msl-2^WT^-FLAG/ Ubi:msl-1^Δ41-85^*); M1[Δ41-65] (♀ *msl-1*^−^*; Ubi:msl-2^WT^-FLAG/ Ubi:msl-1^Δ66-85^*); M1[Δ66-85] (♀ *msl-1*^−^*; Ubi:msl-2^WT^-FLAG/ Ubi:msl-1^Δ66-85^*).

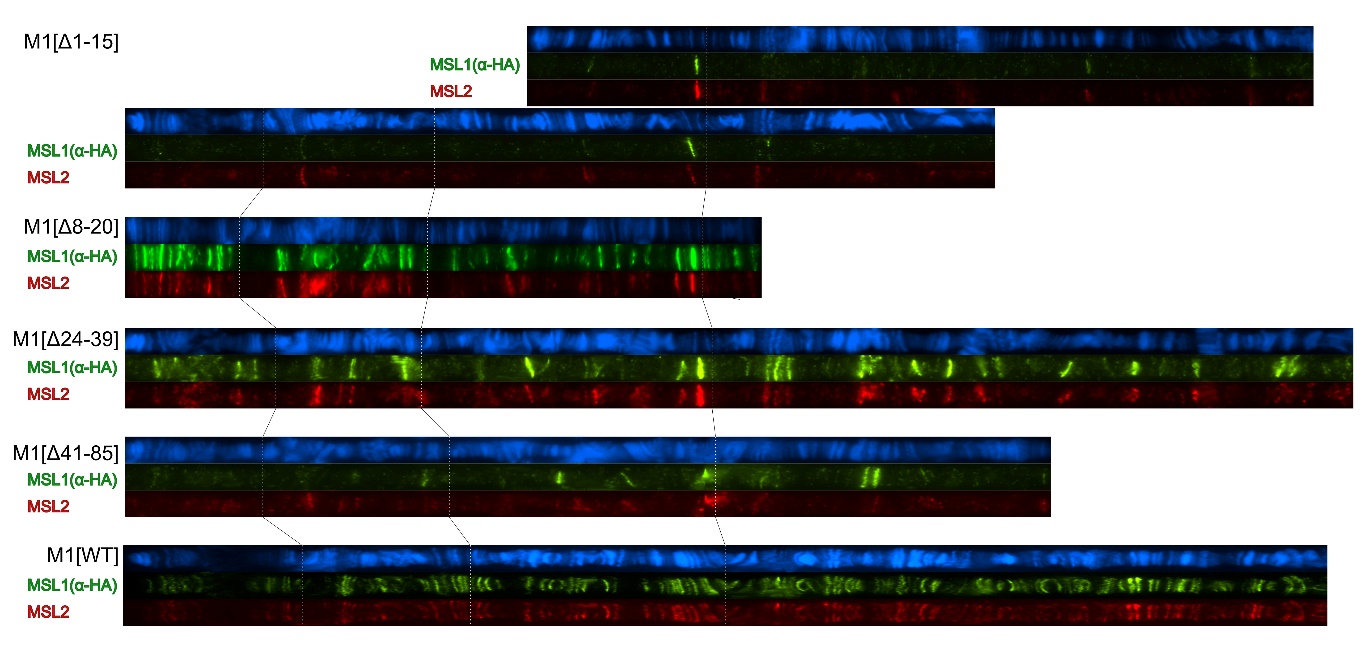

**Figure 2 – figure supplement 2.** Cytological localization of MSL1 variants and MSL2 on the straightened polytene chromosomes of the M1[Δ1-15] (♀ *msl-1*^−^*; Ubi:msl-2^WT^-FLAG/ Ubi:msl-1^Δ1-15^*); M1[Δ8-20] (♀ *msl-1*^−^*; Ubi:msl-2^WT^-FLAG/ Ubi:msl-1^Δ8-20^*); M1[Δ24-39] (♀ *msl-1*^−^*; Ubi:msl-2^WT^-FLAG/ Ubi:msl-1^Δ24-39^*); M1[Δ41-85] (♀ *msl-1*^−^*; Ubi:msl-2^WT^-FLAG/ Ubi:msl-1^Δ41-85^*); M1[WT] (♀ *msl‑1*^−^*; Ubi:msl-2^WT^-FLAG/ Ubi:msl-1^WT^*). Panels show the immunostaining of proteins using mouse anti-HA antibody (green) and rabbit anti-MSL2 antibody (red). DNA was stained with DAPI (blue). Polytene chromosomes were stretched and compared using ImageJ 1.54f/Fiji 2.14.0 software.

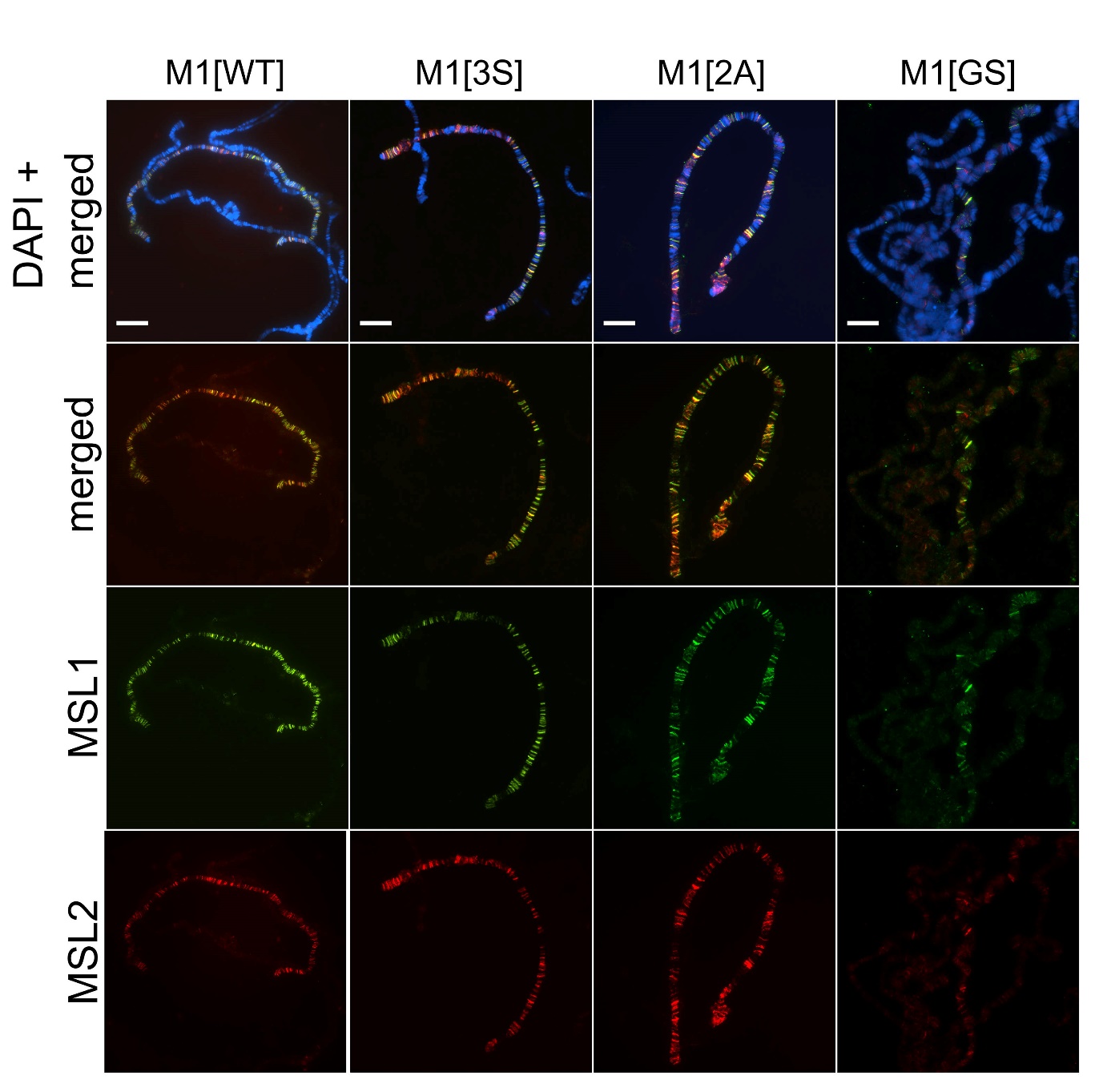

**Figure 3 – figure supplement 1.** Distribution of the MSL complex on the polytene chromosomes from 3rd-day female larvae expressing both MSL1* variants and the MSL2-FLAG protein.

Panels show the results of immunostaining of MSL1 (green, rabbit anti-MSL1 antibody) and MSL2 (red, mouse anti-FLAG antibody). DNA was stained with DAPI (blue). M1[WT] (♀ *msl‑1*^−^*; Ubi:msl-2^WT^-FLAG/ Ubi:msl-1^WT^*); M1[2A] (♀ *msl-1*^−^*; Ubi:msl-2^WT^-FLAG/ Ubi:msl-1^2A^*); M1[3S] (♀ *msl-1*^−^*; Ubi:msl-2^WT^-FLAG/ Ubi:msl-1^3S^*); M1[GS] (♀ *msl-1*^−^*; Ubi:msl-2^WT^-FLAG/ Ubi:msl-1^GS^*).

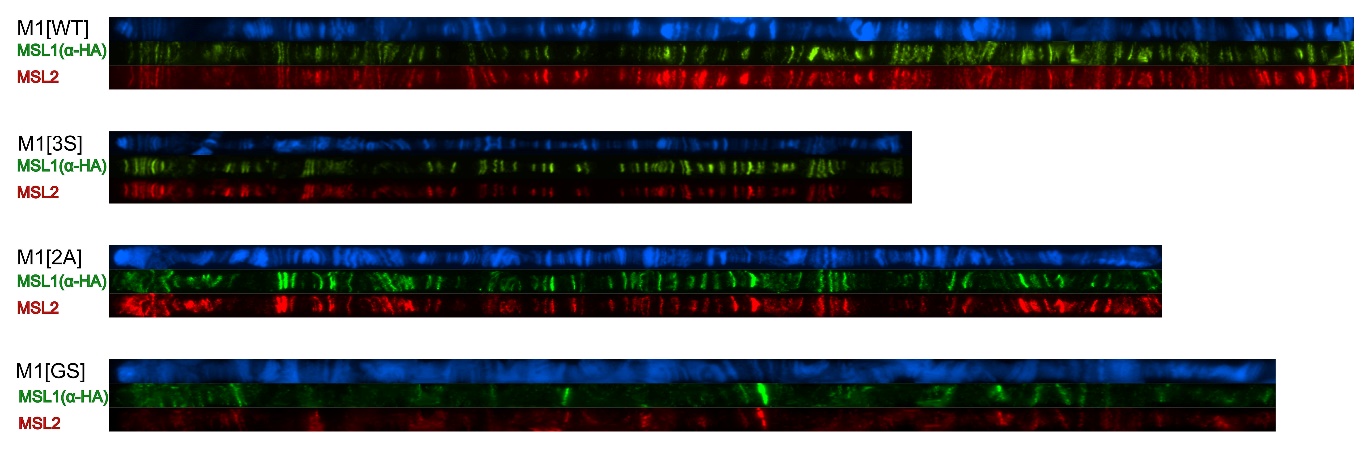

**Figure 3 – figure supplement 2.** Cytological localization of MSL1 variants and MSL2 on the straightened polytene chromosomes of the M1[WT] (♀ *msl-1*^−^*; Ubi:msl-2^WT^-FLAG/ Ubi:msl-1^WT^*); M1[2A] (♀ *msl-1*^−^*; Ubi:msl-2^WT^-FLAG/ Ubi:msl-1^2A^*); M1[3S] (♀ *msl-1*^−^*; Ubi:msl-2^WT^-FLAG/ Ubi:msl-1^3S^*); M1[GS] (♀ *msl-1*^−^*; Ubi:msl-2^WT^-FLAG/ Ubi:msl-1^GS^*). Panels show the immunostaining of proteins using mouse anti-HA antibody (green) and rabbit anti-MSL2 antibody (red). DNA was stained with DAPI (blue).

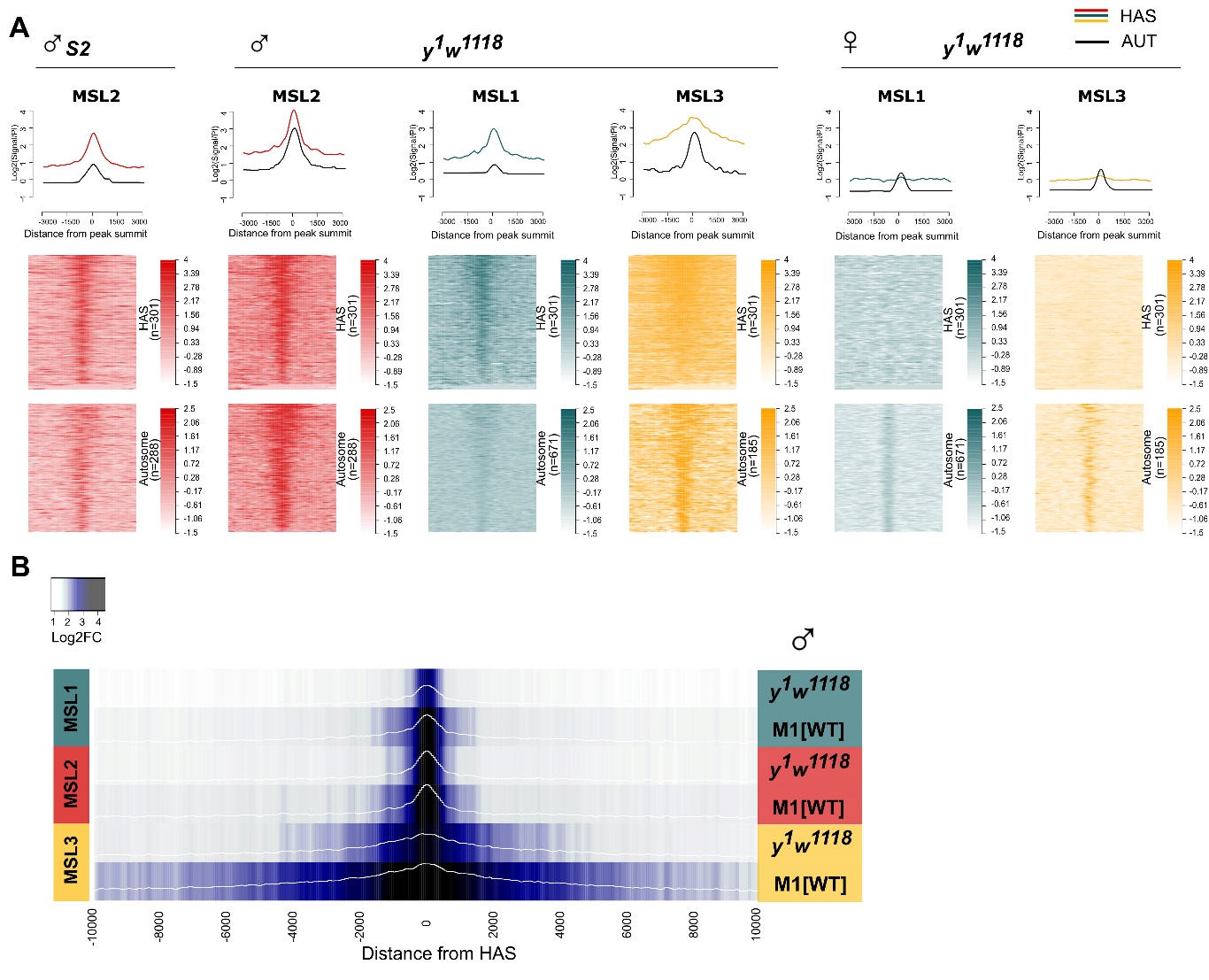

**Figure 5 – figure supplement 1.** **(A)** Comparison of MSL1 (green), MSL2 (red), MSL3 (yellow) binding in 2–3-day-old males (♂) and females (♀) of the *y^1^w^1118^* line at HAS and autosomal sites for MSL proteins. Average signal (on the top) and heatmap (on the bottom) showing the log-fold change (FC) between normalized (RPKM) test signals and nonspecific IgG signals in HAS and autosomal regions (see Materials and Methods). MSL2 S2 track was obtained from (Schauer et al., 2017). LogFC was calculated after smoothing signals using Daniell kernel, with kernel size 50 and the addition of a pseudocount. On the heatmaps, the peaks are ranked according to the average logFC in the ♂M1[wt] sample. **(B)** Comparing of MSL1, MSL2 and MSL3 binding to HAS in 2–3-day-old *y^1^w^1118^* and M1[wt] (*msl-1*^−^*; Ubi:msl-1^WT^* */ Ubi:msl-1^WT^* ) males. Average log fold-change between normalized (RPKM) test signals and nonspecific IgG signals in HAS (see Materials and Methods). Average log fold-change was calculated after smoothing signals using the Daniell kernel with kernel size 50 and the addition of a pseudocount. Next, the tracks were aggregated using 100-bp bins.

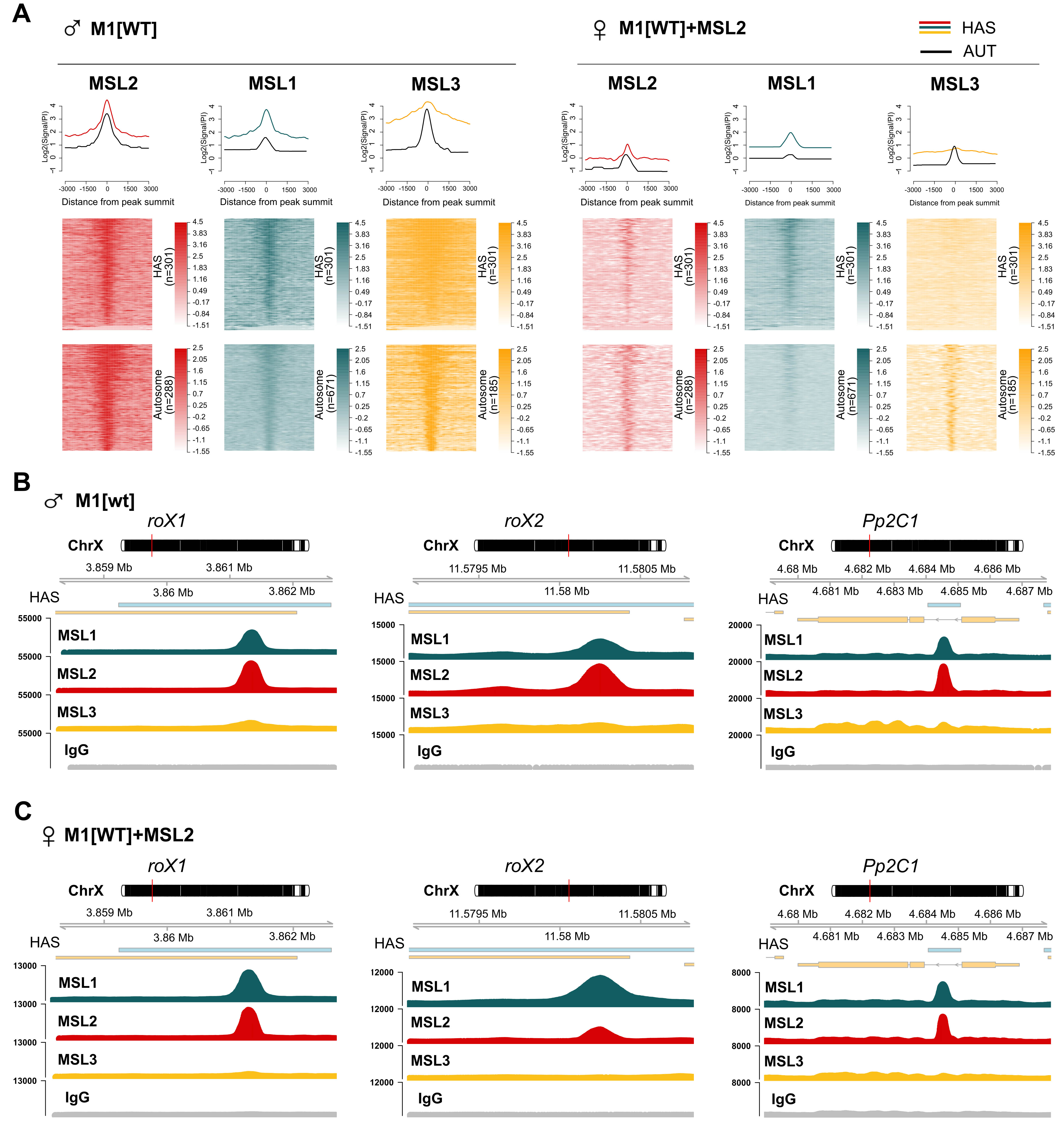

**Figure 5 – figure supplement 2.** Comparisons of the binding patterns of MSL proteins between 2-3-day adult males and females expressing MSL2. (A) Average signal (top) and heatmap (bottom) showing the log fold-change (FC) between normalized (RPKM) test signals and nonspecific IgG signals for HAS and autosomal regions (see Materials and Methods). LogFC was calculated after smoothing signals using the Daniell kernel with kernel size 50 and the addition of a pseudocount. On the heatmaps, the peaks are ranked according to the average logFC in the M1[wt] male sample. ♂ M1[wt] indicates 2–3-day-old *msl-1*^−^*; Ubi:msl-1^WT^/ Ubi:msl-1^WT^* males. ♀M1[wt]+MSL2 indicates 2–3-day-old *msl-1*^−^*; Ubi:msl-2* *^WT^-FLAG/ Ubi:msl-1^WT^* females. (B and C) Enrichment of MSL1 (green), MSL2 (red), and MSL3 (yellow) signals associated with the X-linked genes encoding *Pp2C1* and the non-coding RNAs *roX1* and *roX2* in (B) M1[WT] males and (C) M1[WT]+MSL2 females.

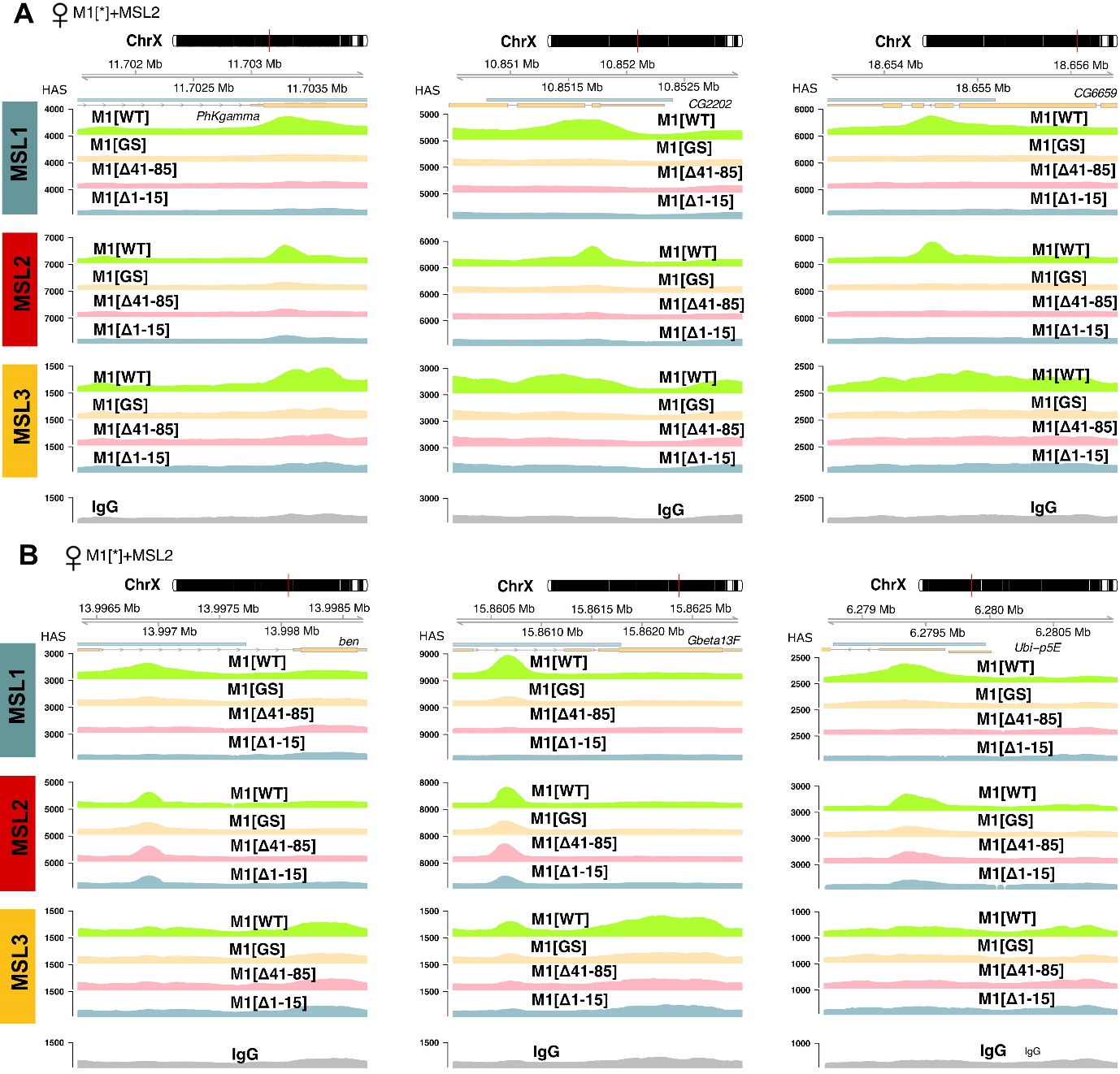

**Figure 5 – figure supplement 3.** Examples of MSL1, MSL2, and MSL3 signal enrichment associated with X chromosomal HAS in M1[WT] (♀ *msl-1*^−^*; Ubi:msl-2^WT^-FLAG/ Ubi:msl-1^WT^*), M1[GS] (♀ *msl-1*^−^*; Ubi:msl-2^WT^-FLAG/ Ubi:msl-1^GS^*), M1[Δ41-85] (♀ *msl-1*^−^*; Ubi:msl-2^WT^-FLAG/ Ubi:msl-1^Δ41-85^*), M1[Δ1-15] (♀ *msl-1*^−^*; Ubi:msl-2^WT^-FLAG/ Ubi:msl-1^Δ1-15^*) females. MSL2 binds to HAS **(A)** only in M1[WT] females or **(B)** in M1[WT], M1[GS], M1[Δ41-85] and M1[Δ1-15] females.

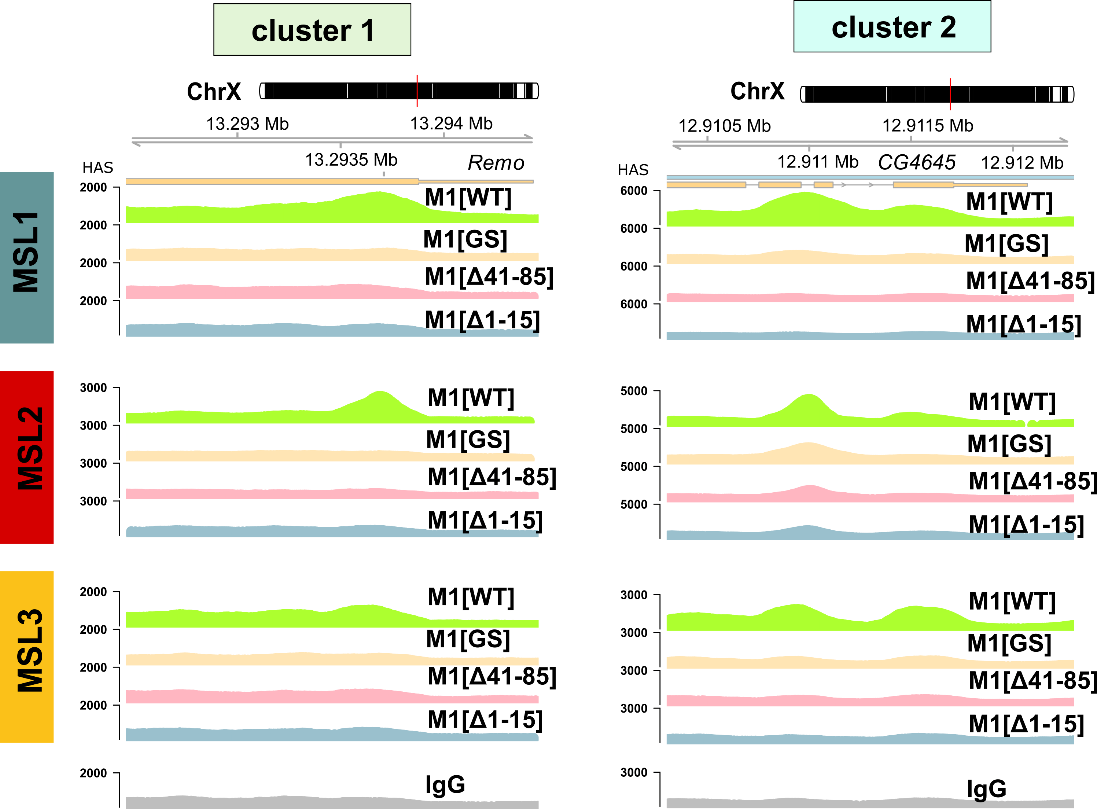

**Figure 6 – figure supplement 1.** Examples of MSL1, MSL2, and MSL3 signal enrichment in M1[WT] (♀ *msl-1*^−^*; Ubi:msl-2^WT^-FLAG/ Ubi:msl-1^WT^*), M1[GS] (♀ *msl-1*^−^*; Ubi:msl-2^WT^-FLAG/ Ubi:msl-1^GS^*), M1[Δ41-85] (♀ *msl-1*^−^*; Ubi:msl-2^WT^-FLAG/ Ubi:msl-1^Δ41-85^*), M1[Δ1-15] (♀ *msl-1*^−^*; Ubi:msl-2^WT^-FLAG/ Ubi:msl-1^Δ1-15^*) females. Clusters 1 and 2 are associated with MSL2 binding to X chromosomal sites.

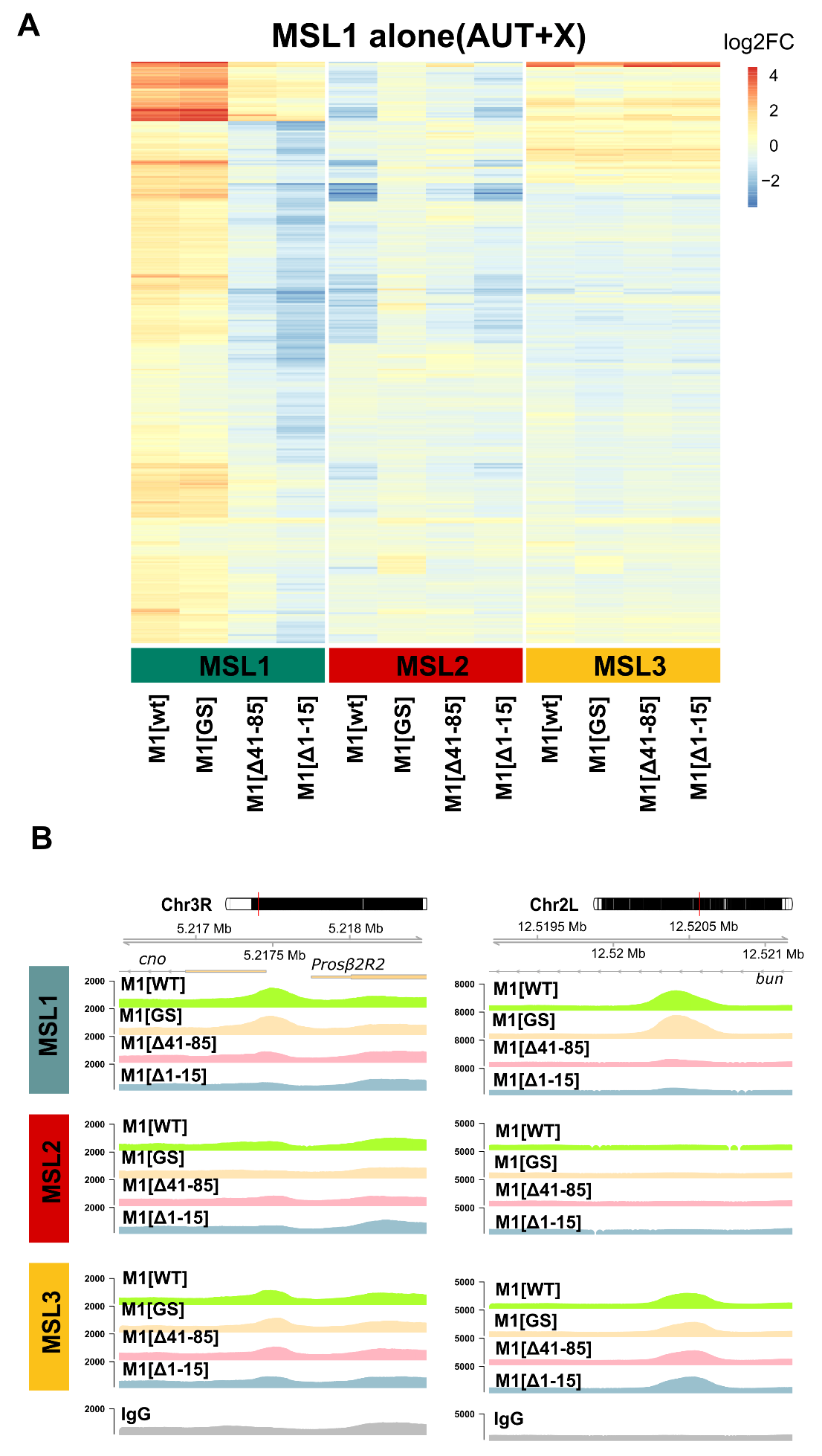

**Figure 6 – figure supplement 2.** Regions with MSL1 alone (no colocalization with MSL2) peaks (autosomes and X chromosome). **(A)** Heatmap of average logFC between MSL and nonspecific IgG signal in female fly lines. **(B)** Examples of MSL1, MSL2, and MSL3 signal enrichment at autosomes in M1[WT] (♀ *msl-1*^−^*; Ubi:msl-2^WT^-FLAG/ Ubi:msl-1^WT^*), M1[GS] (♀ *msl-1*^−^*; Ubi:msl-2^WT^-FLAG/ Ubi:msl-1^GS^*), M1[Δ41-85] (♀ *msl-1*^−^*; Ubi:msl-2^WT^-FLAG/ Ubi:msl-1^Δ41-85^*), M1[Δ1-15] (♀ *msl-1*^−^*; Ubi:msl-2^WT^-FLAG/ Ubi:msl-1^Δ1-15^*) females.
