## Supplementary figures and images for "N-Terminus of *Drosophila Melanogaster* MSL1 Is Critical for Dosage Compensation"

### A_ lamin.tif

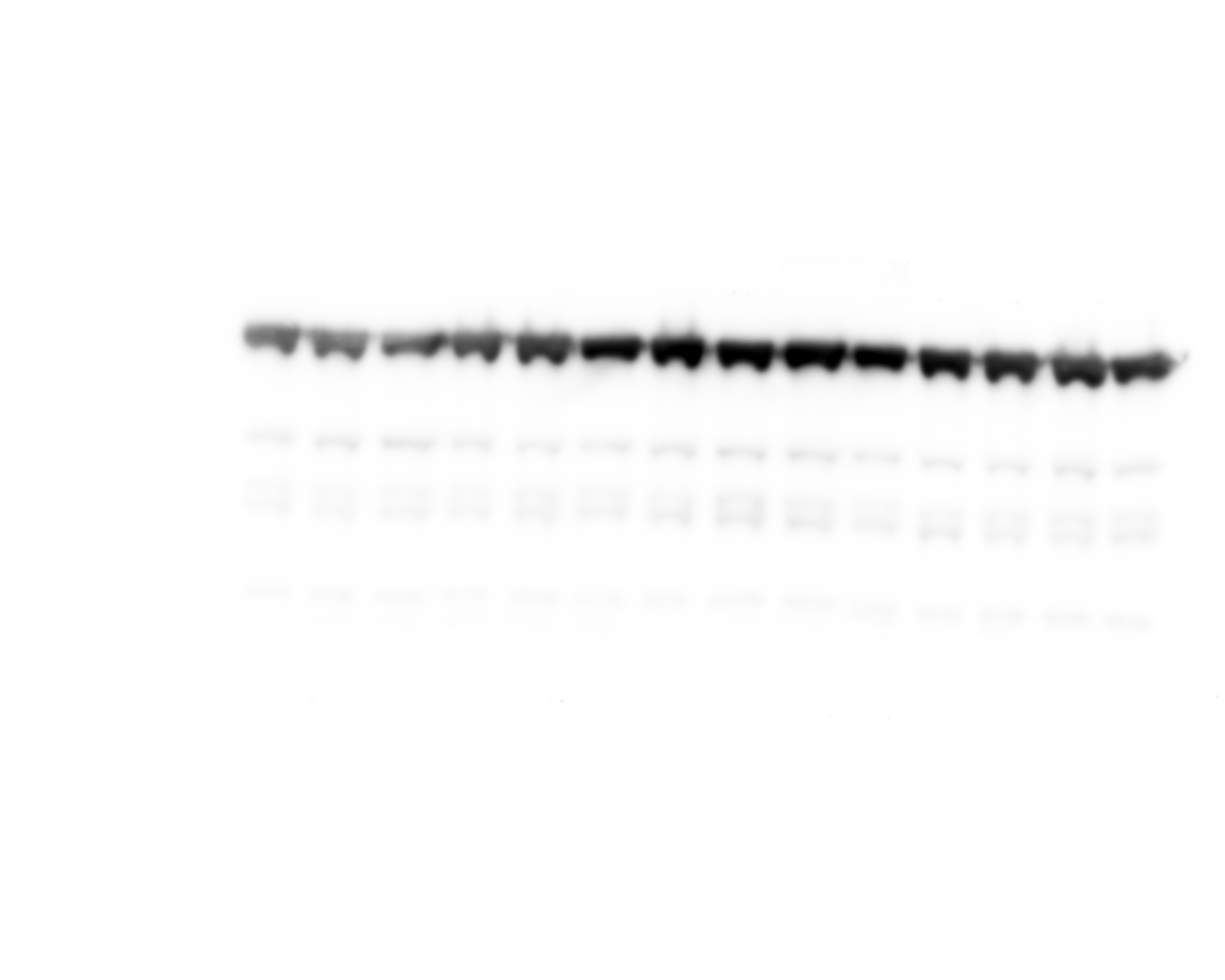

### A_FLAG msl2 S2.tif

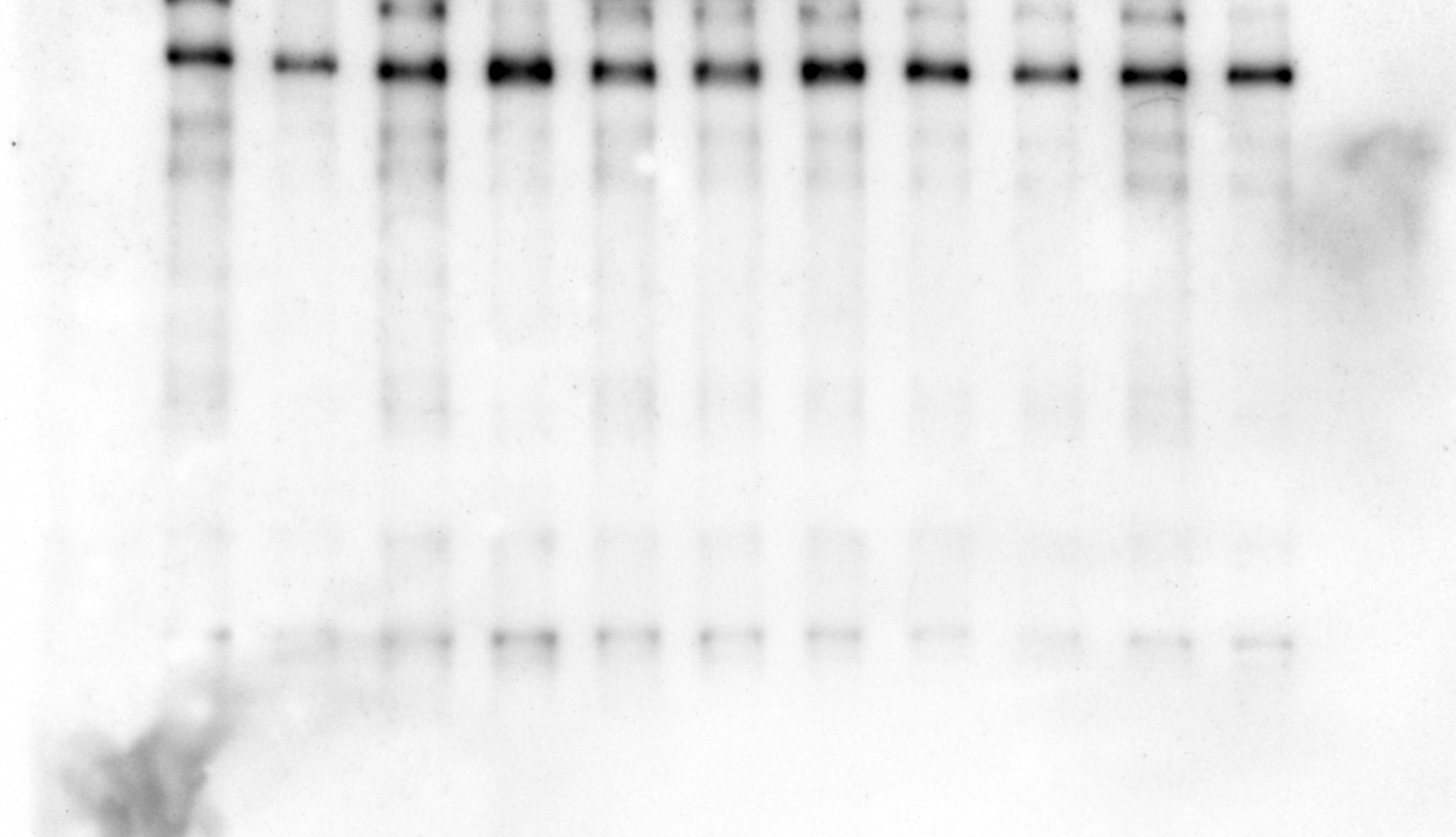

### A_HA msl1 S2.tif

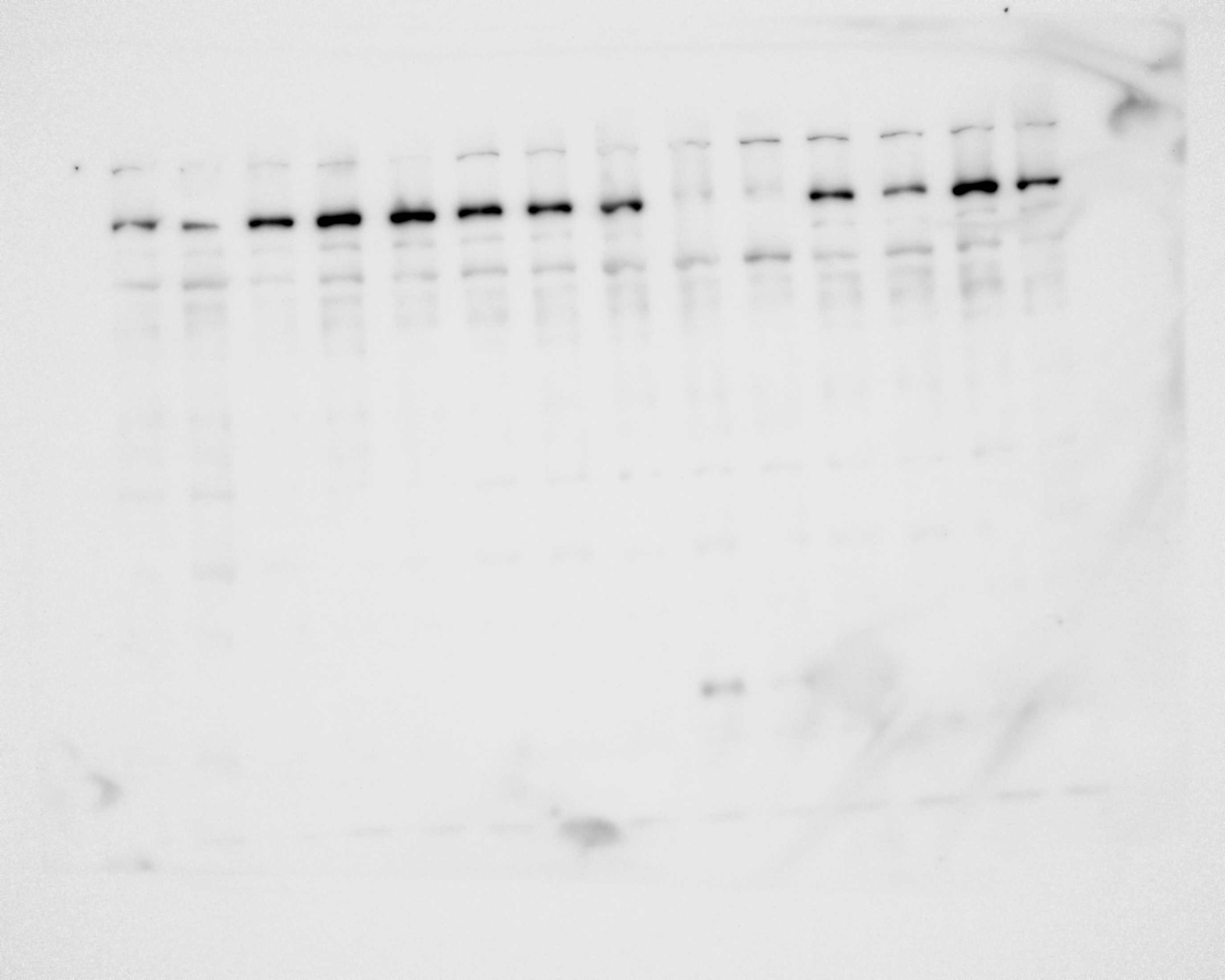

### B _ msl2(Composite).tif

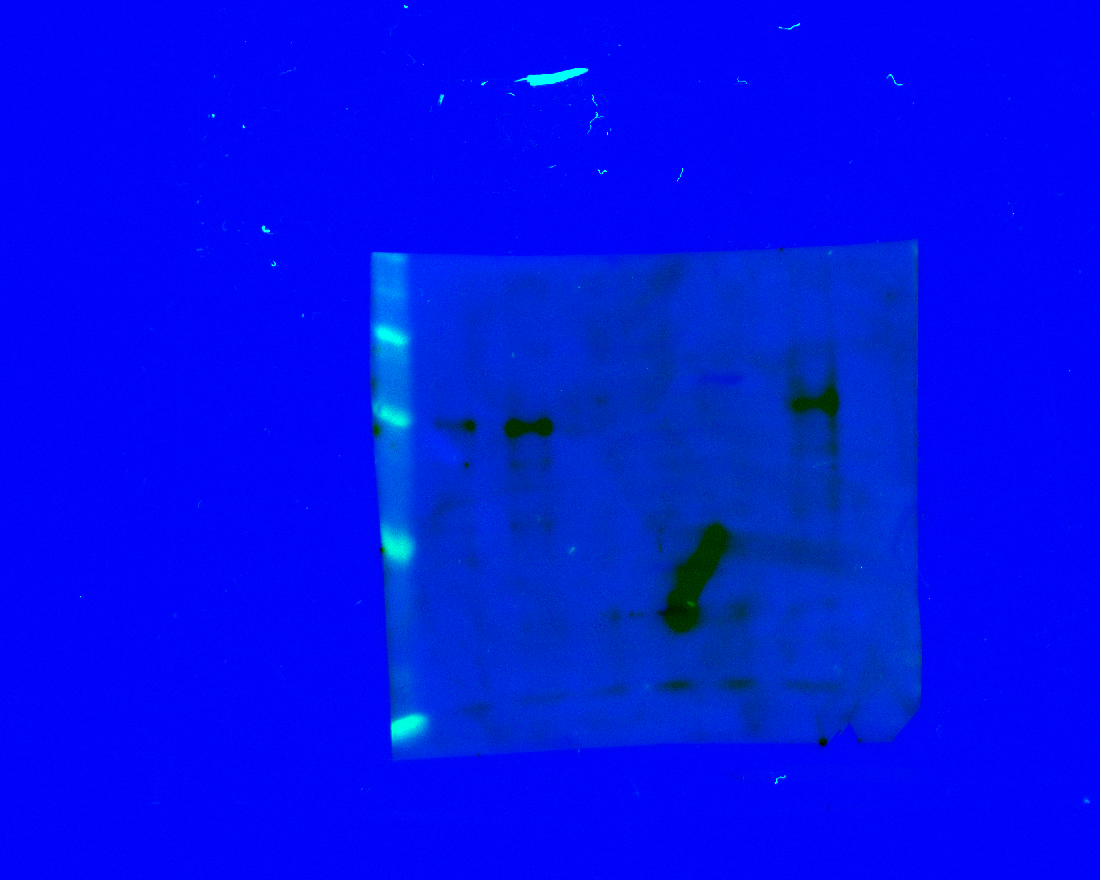

### B _ msl2.tif

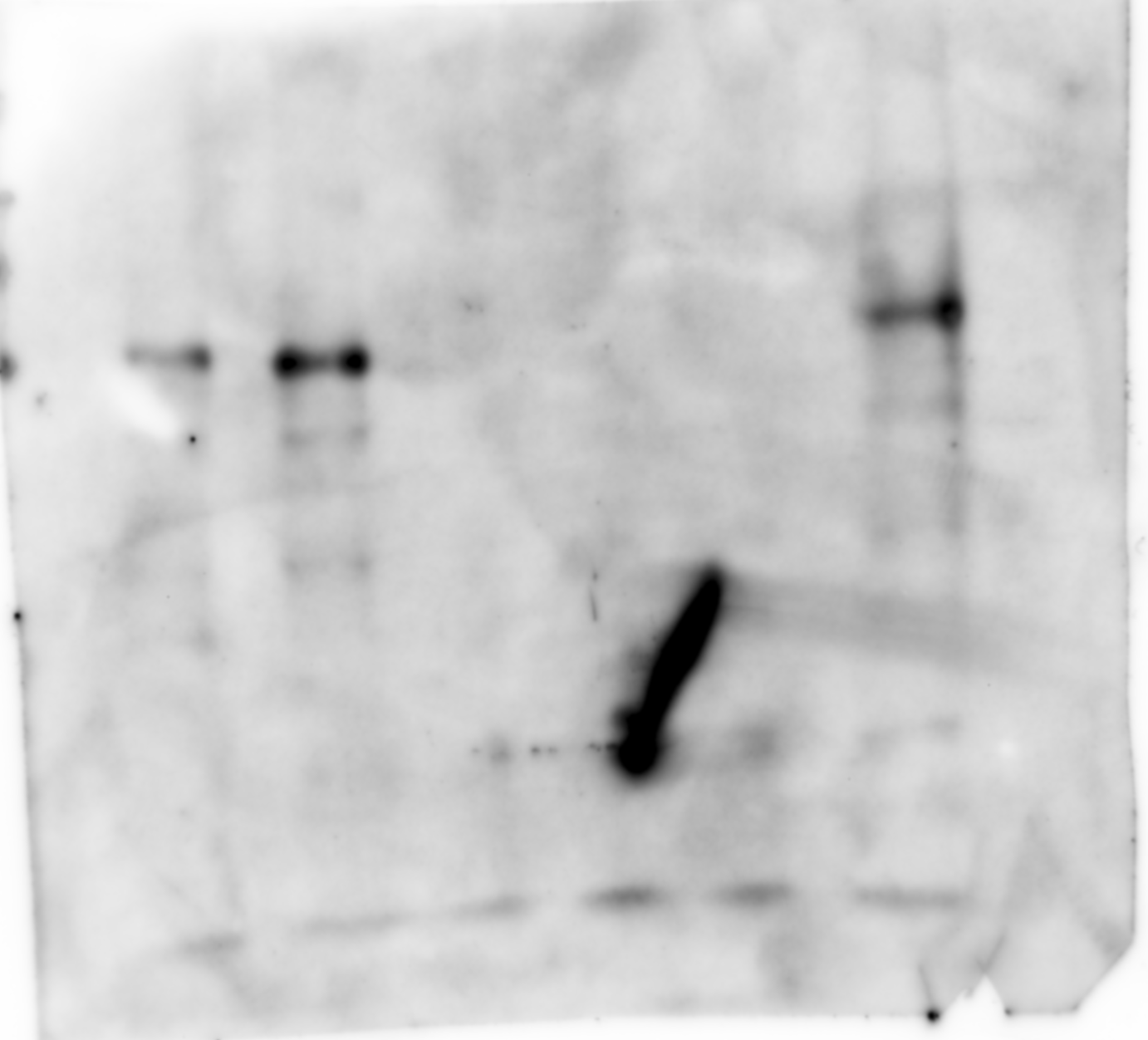

### B_ lamin(Composite).tif

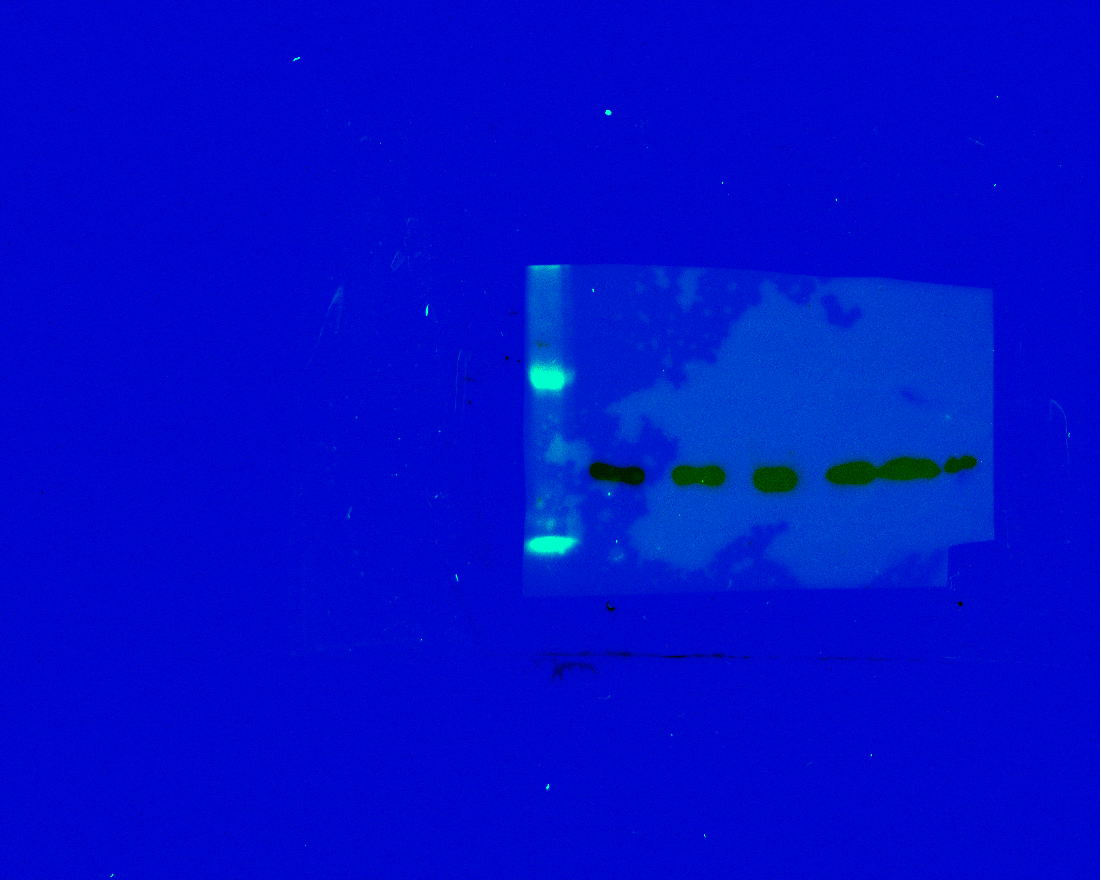

### B_ lamin.tif

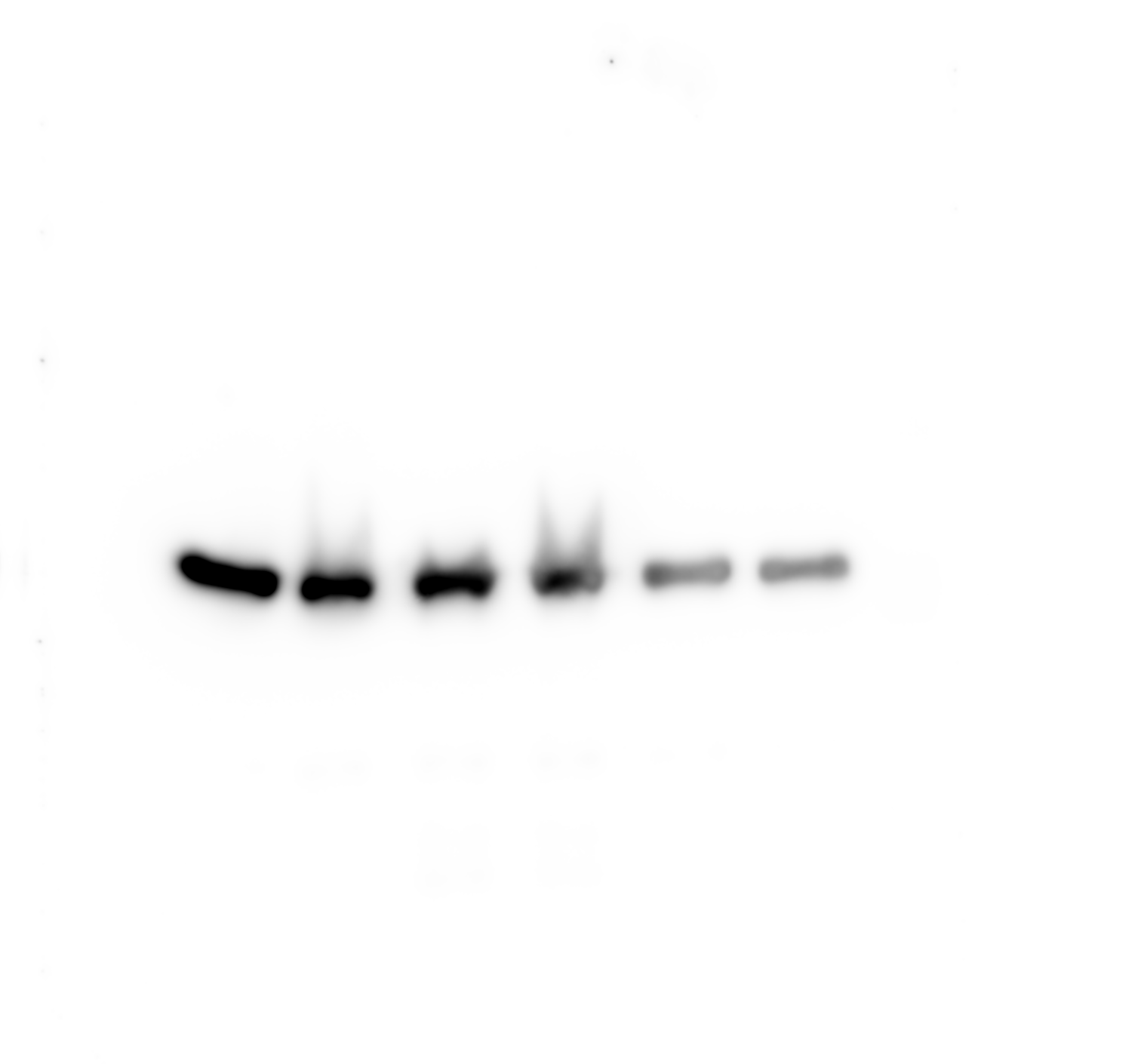

### B_ msl1(Composite).tif

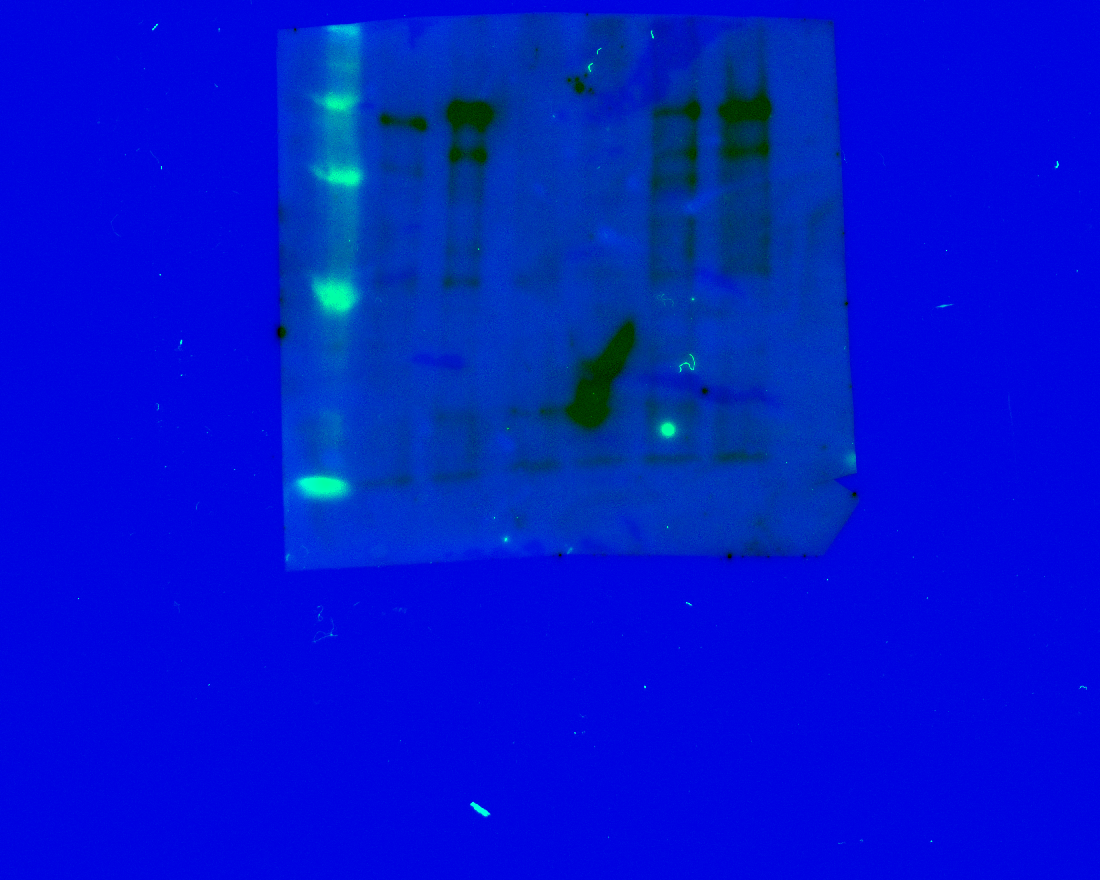

### B_ msl1.tif

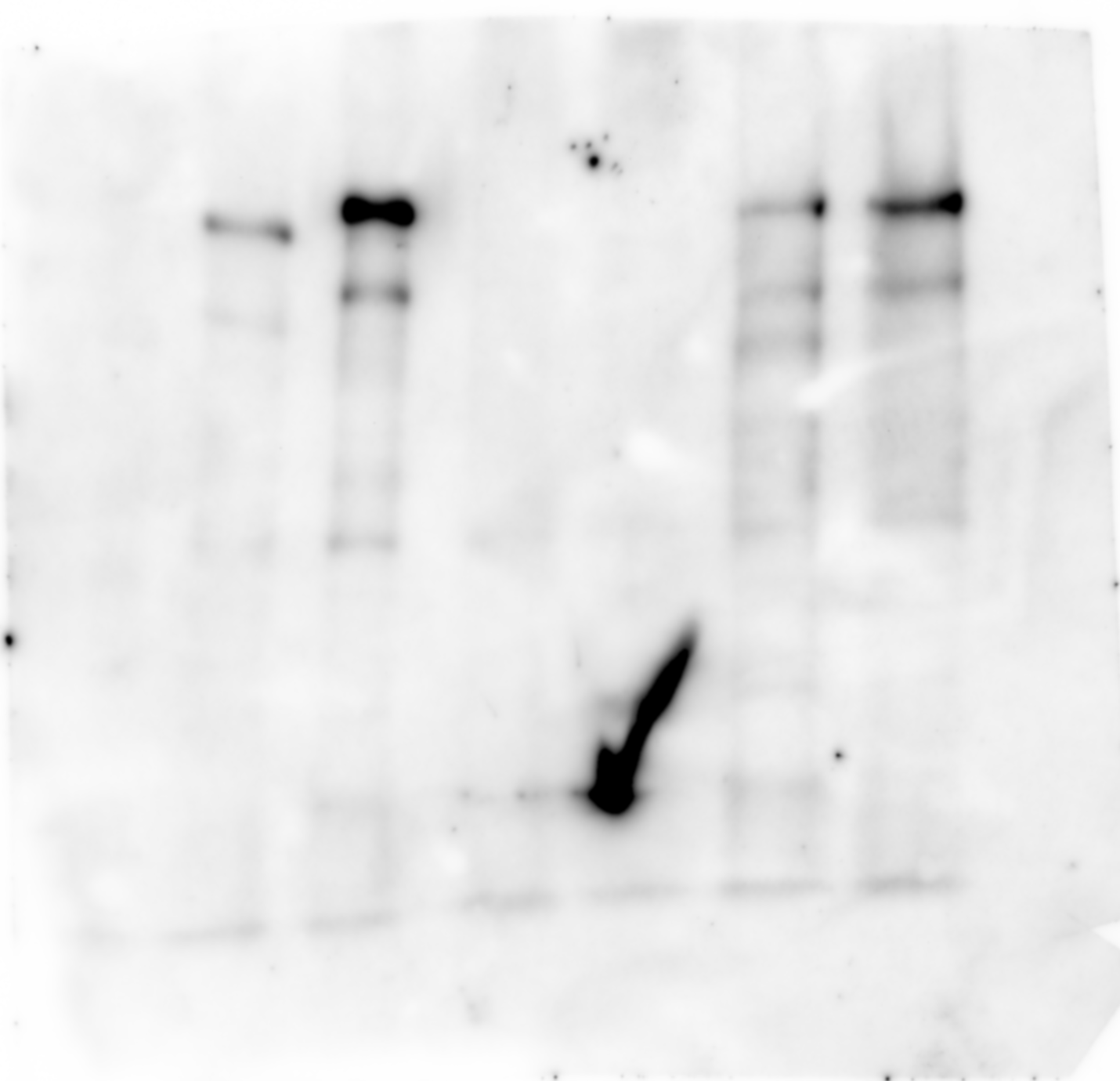

### B_flag ip 2mut gs(Chemiluminescence).tif

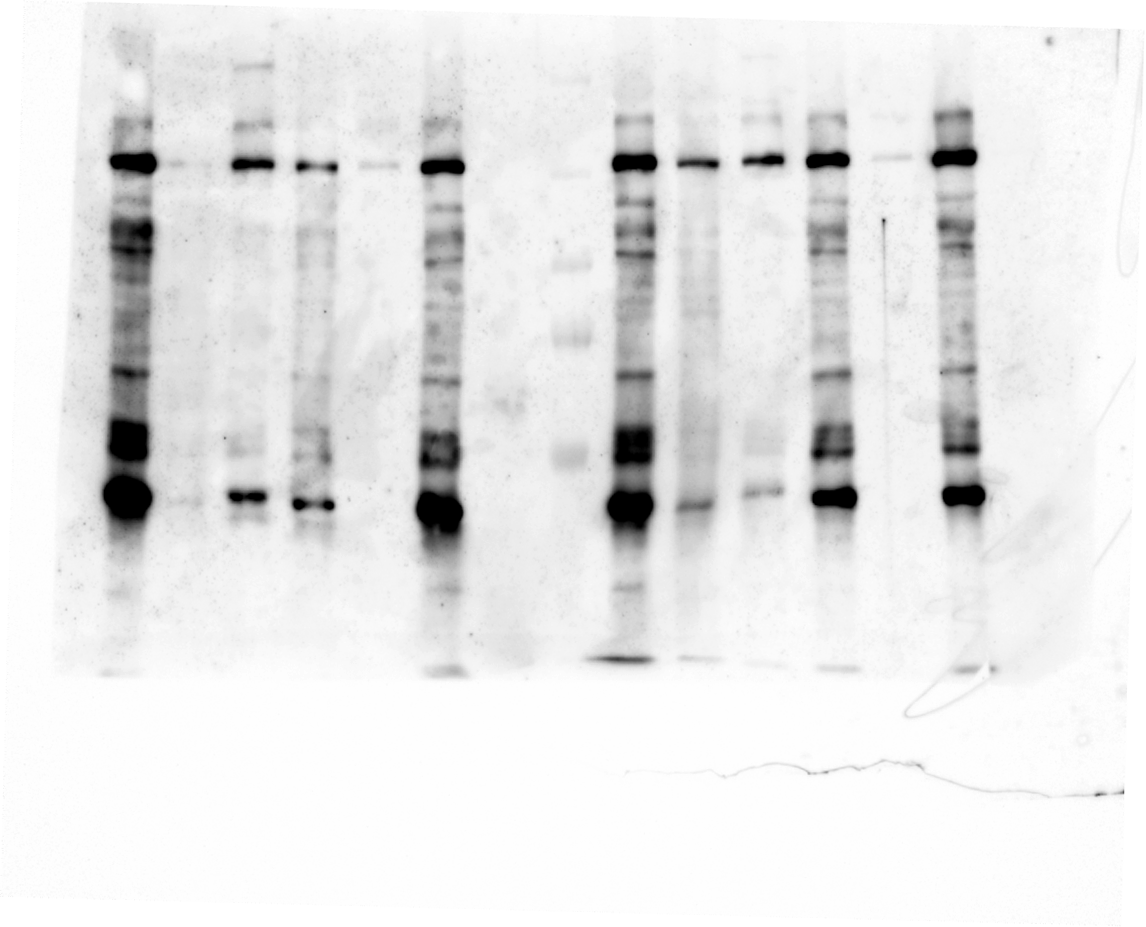

### B_flag ip 3mut 3mutA 2(Chemiluminescence).tif

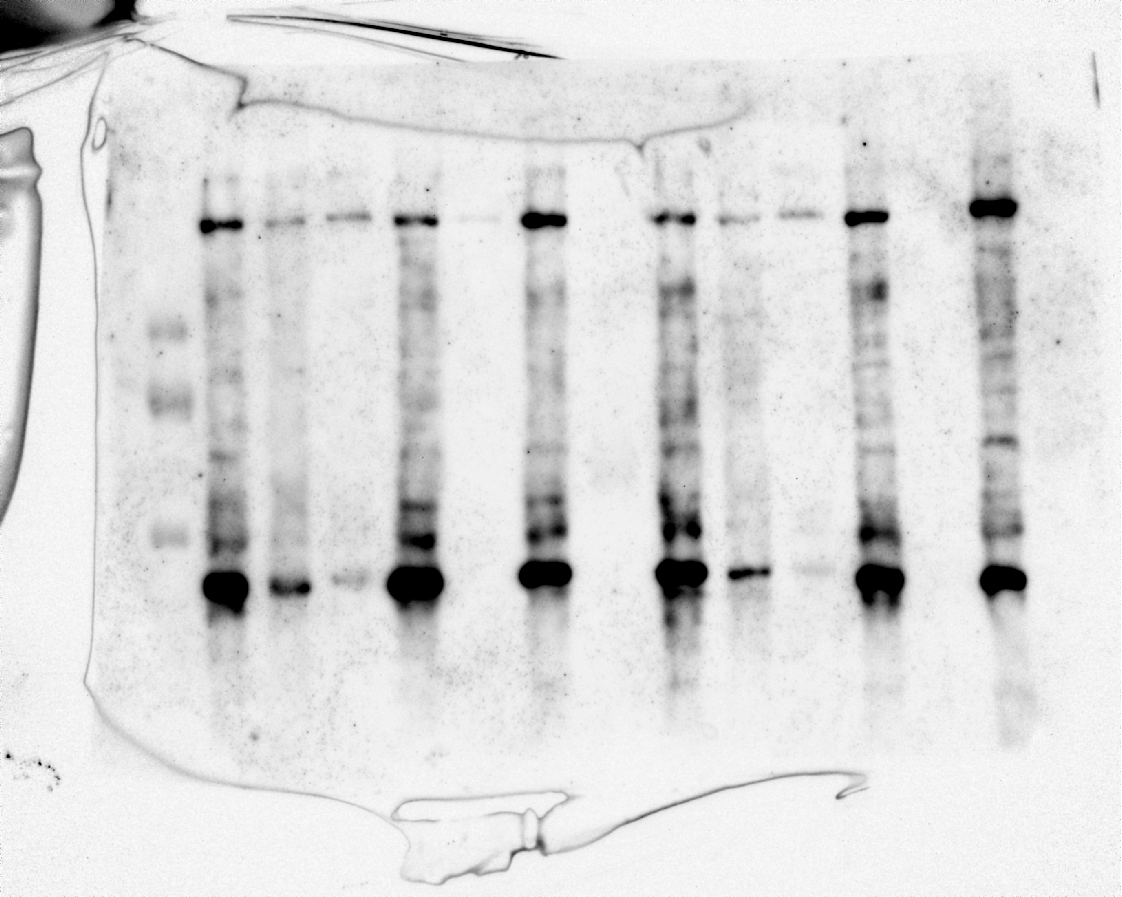

### B_flag ip 3mut gs(Chemiluminescence).tif

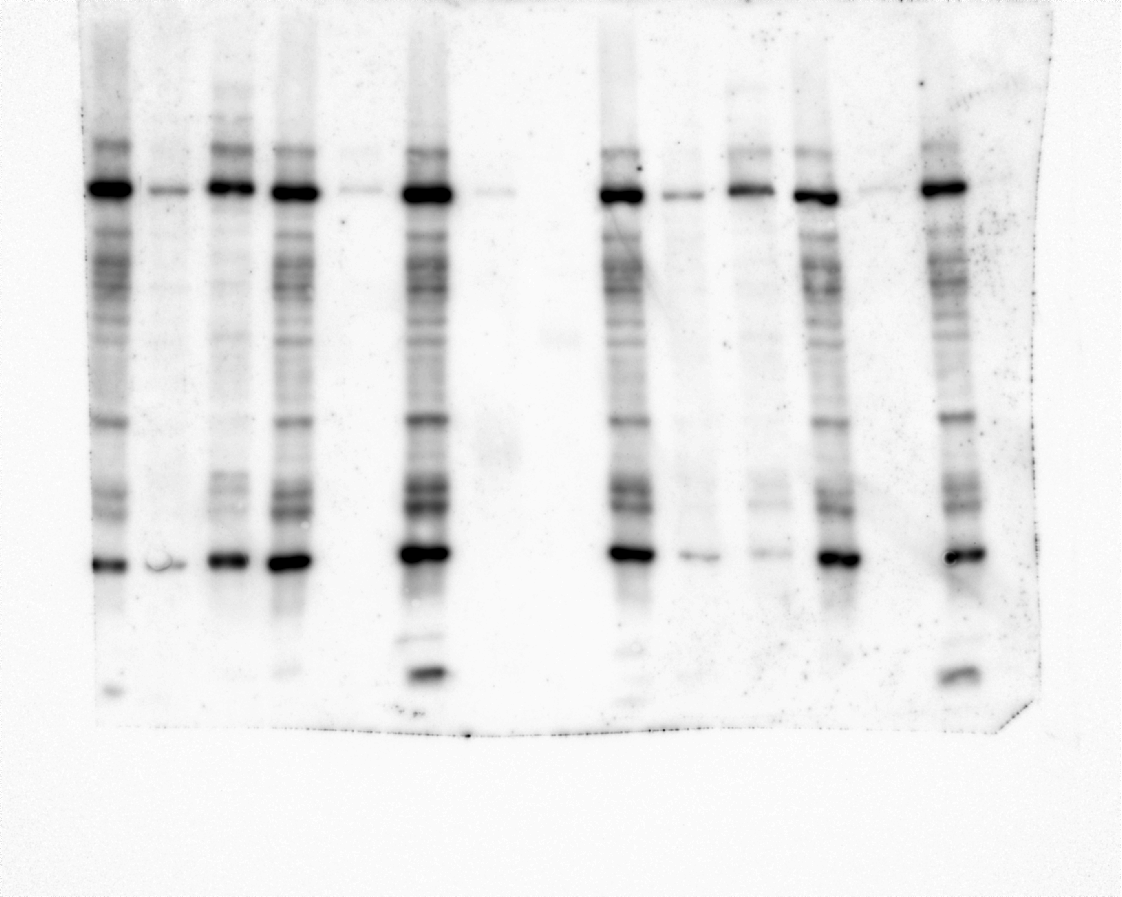

### B_flag ip 8-20 24-39(Chemiluminescence).tif

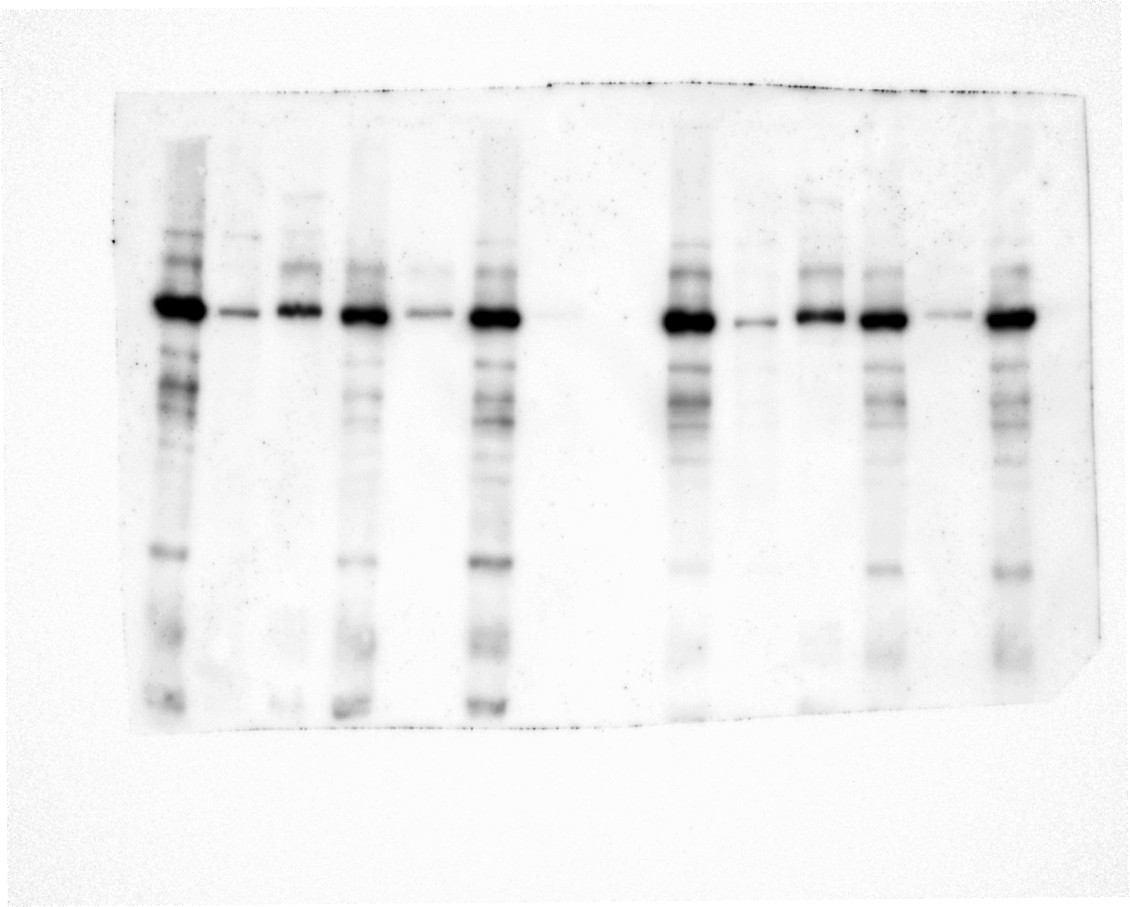

### B_flag ip 24-39 8-20(Chemiluminescence).tif

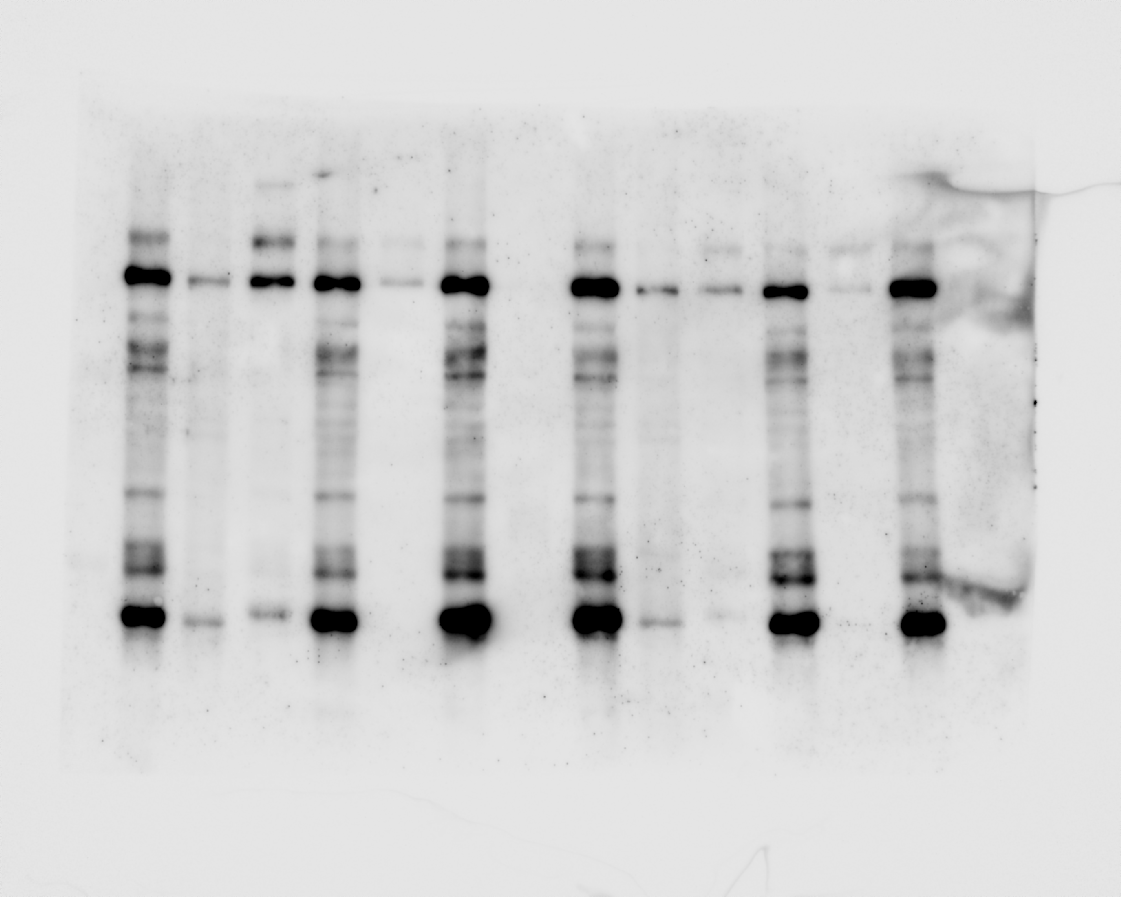

### B_flag ip 41-85(Chemiluminescence).tif

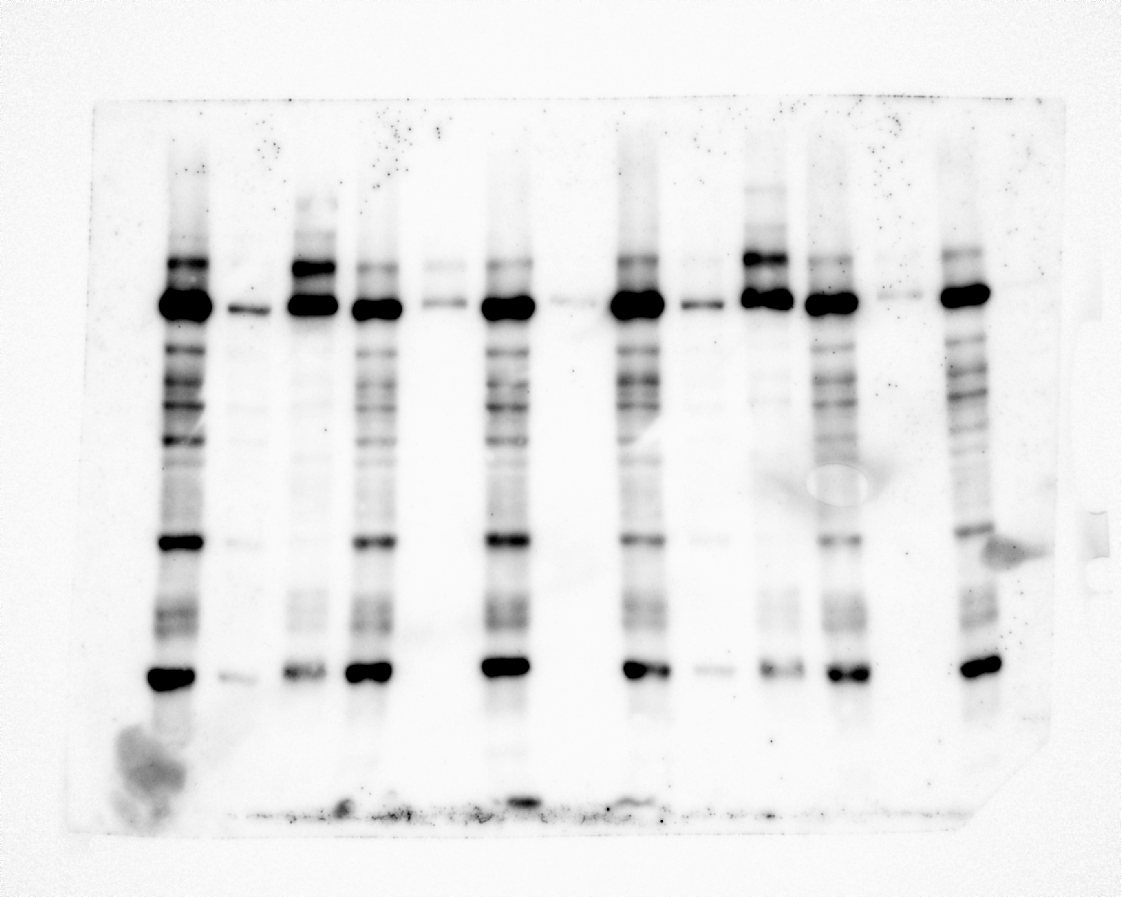

### B_flag ip wt 3mut_2(Chemiluminescence).tif

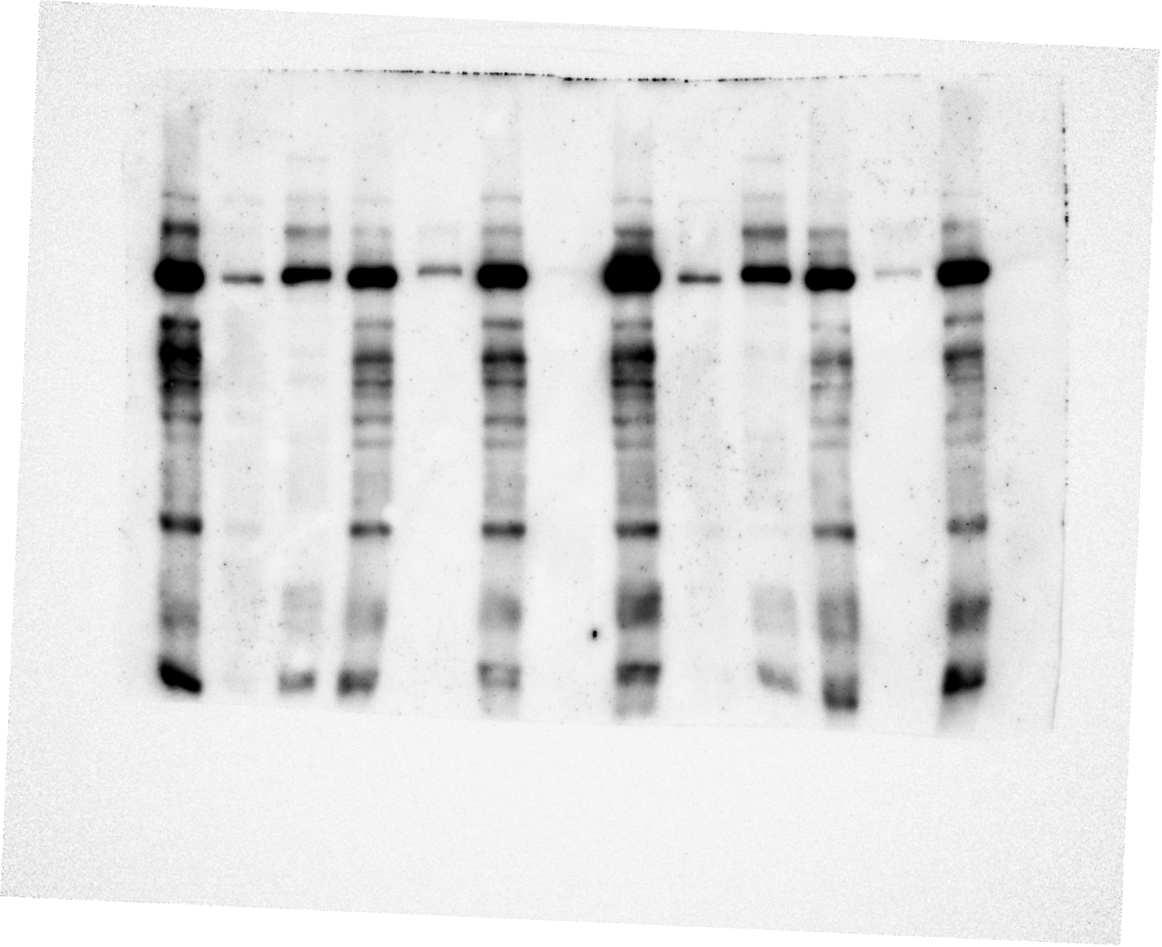

### B_flag ip wt(Chemiluminescence).tif

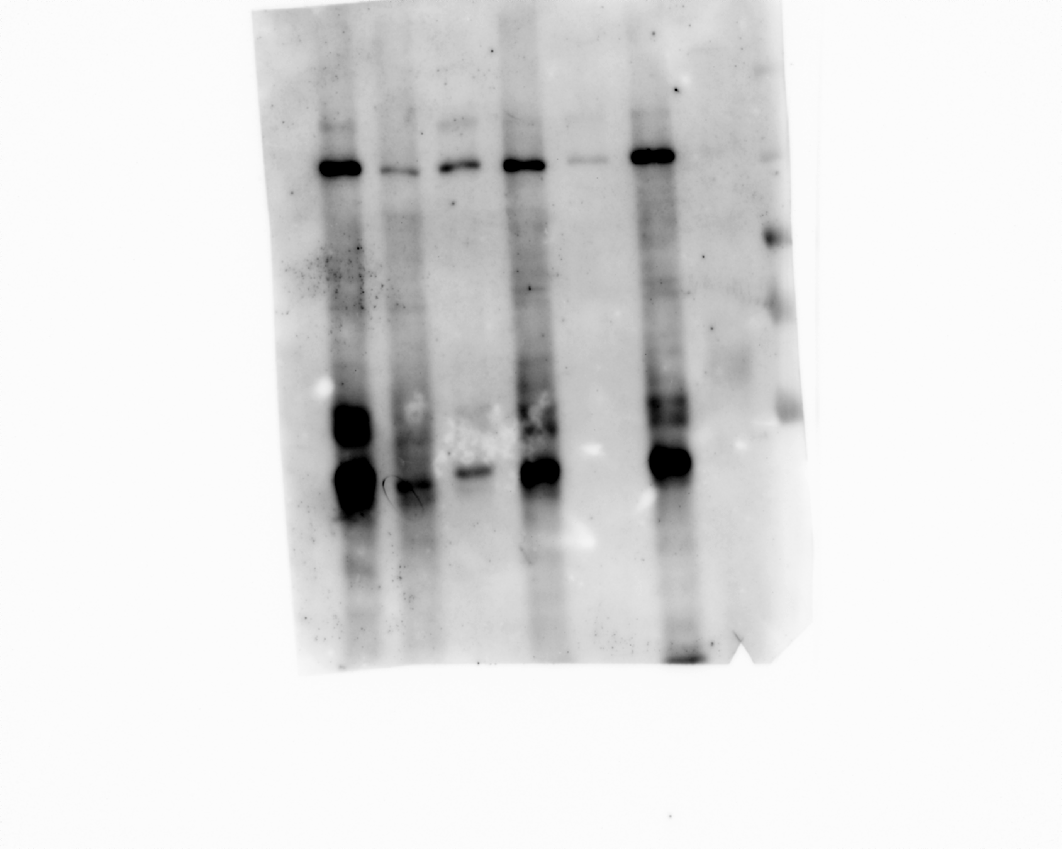

### B_ip msl1_ha_6(Chemiluminescence).raw16.tif

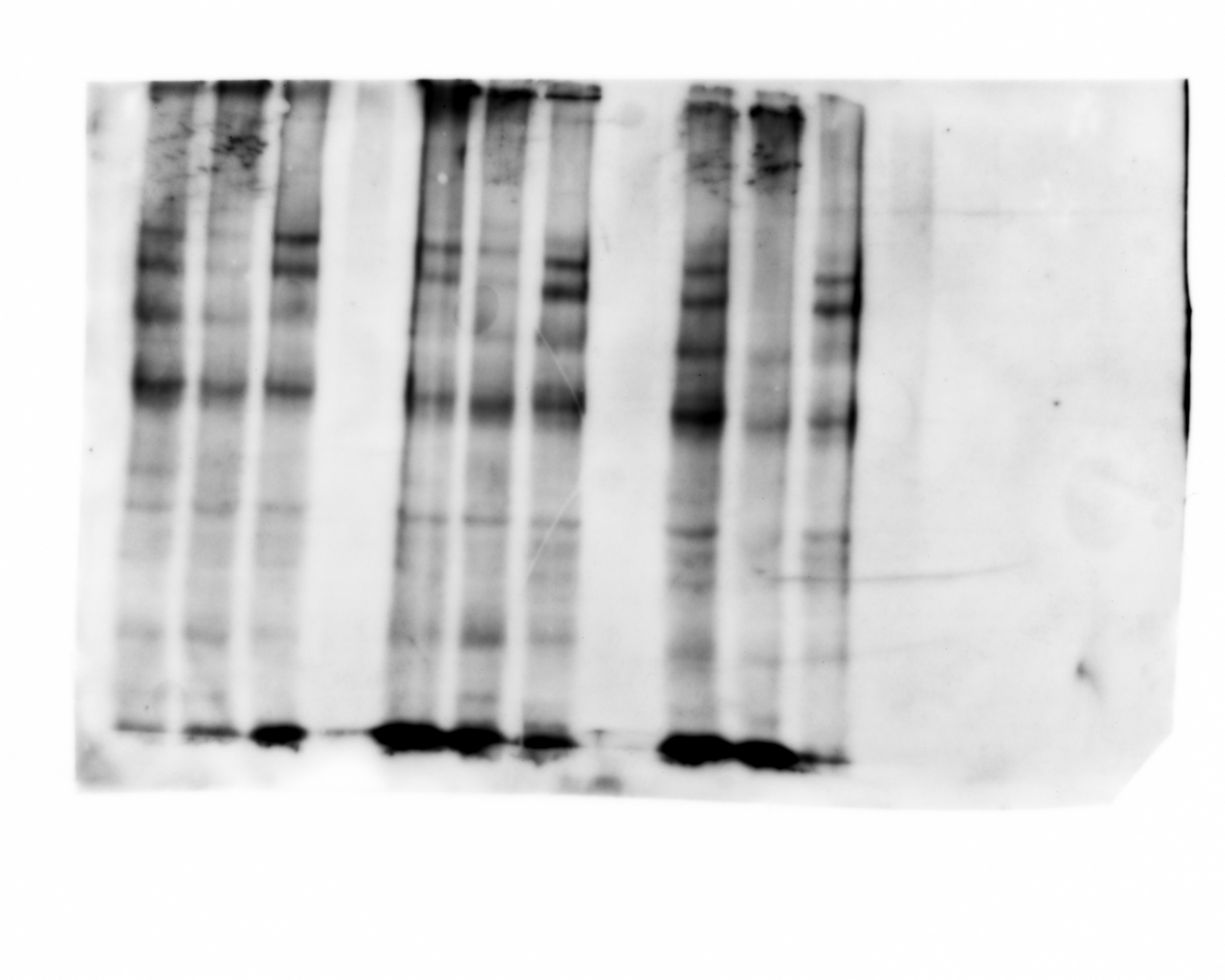

### B_lamin_m1_wtgs2a3s_4.tif

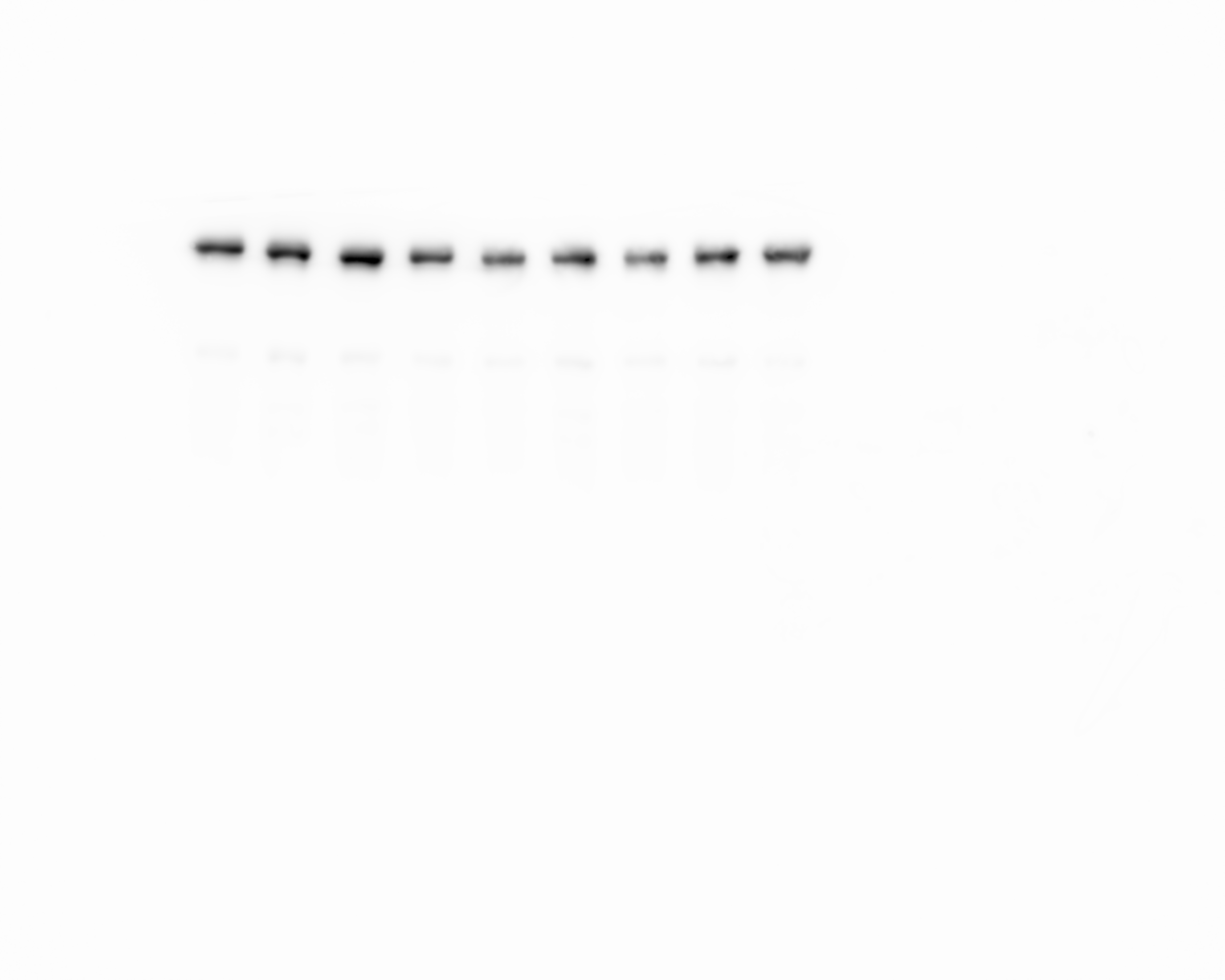

### BC_msl1_wt2a3sgs.tif

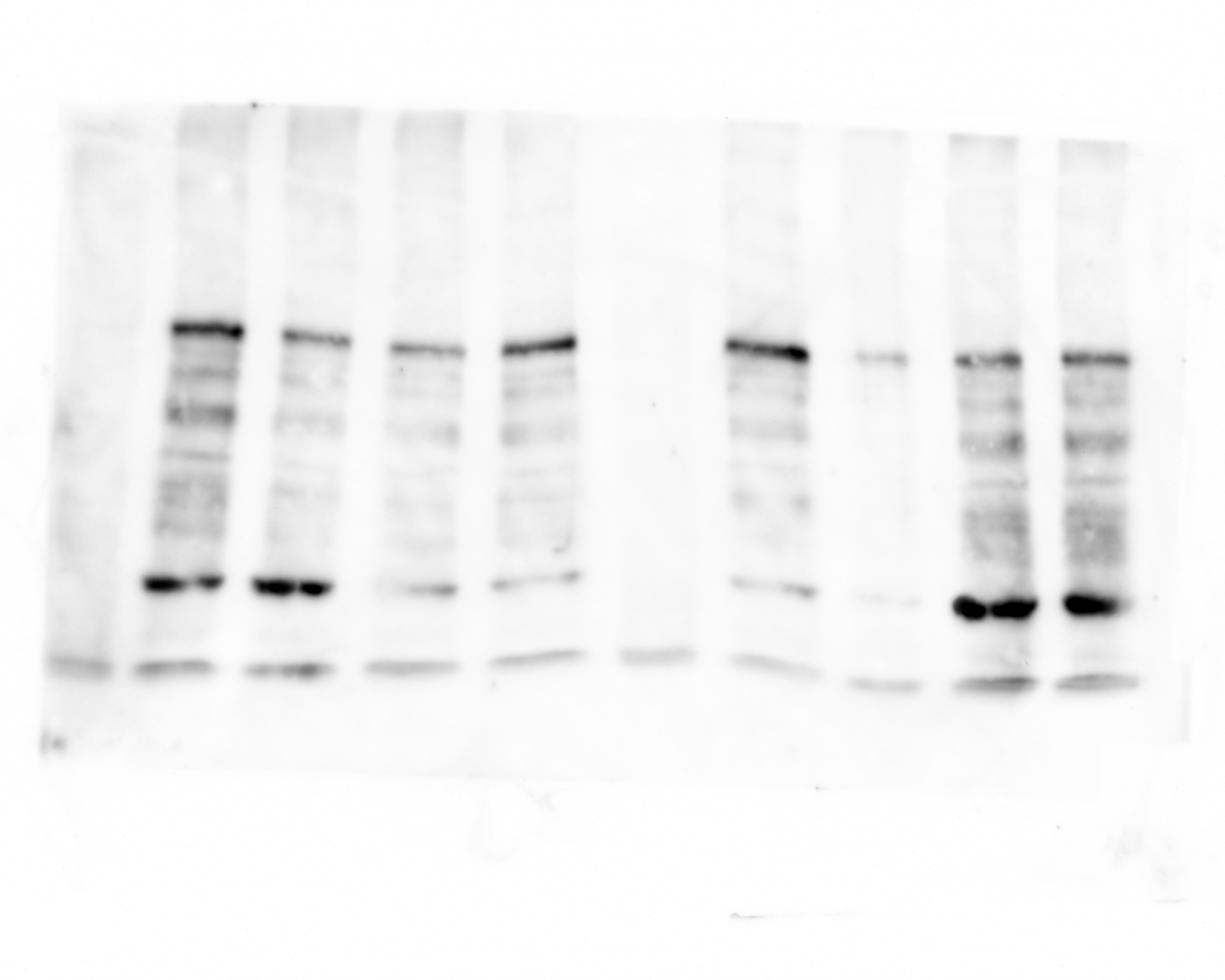

### C_ lamin.tif

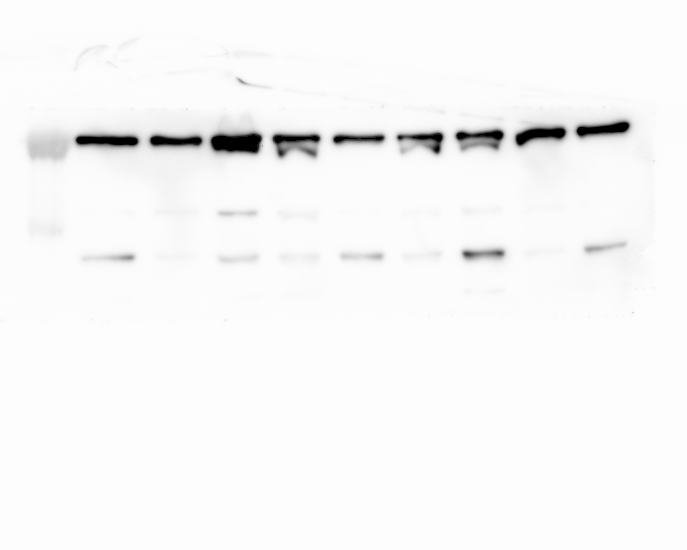

### C_lamin_m1m2_wtgs2a3s_4(Composite).tif

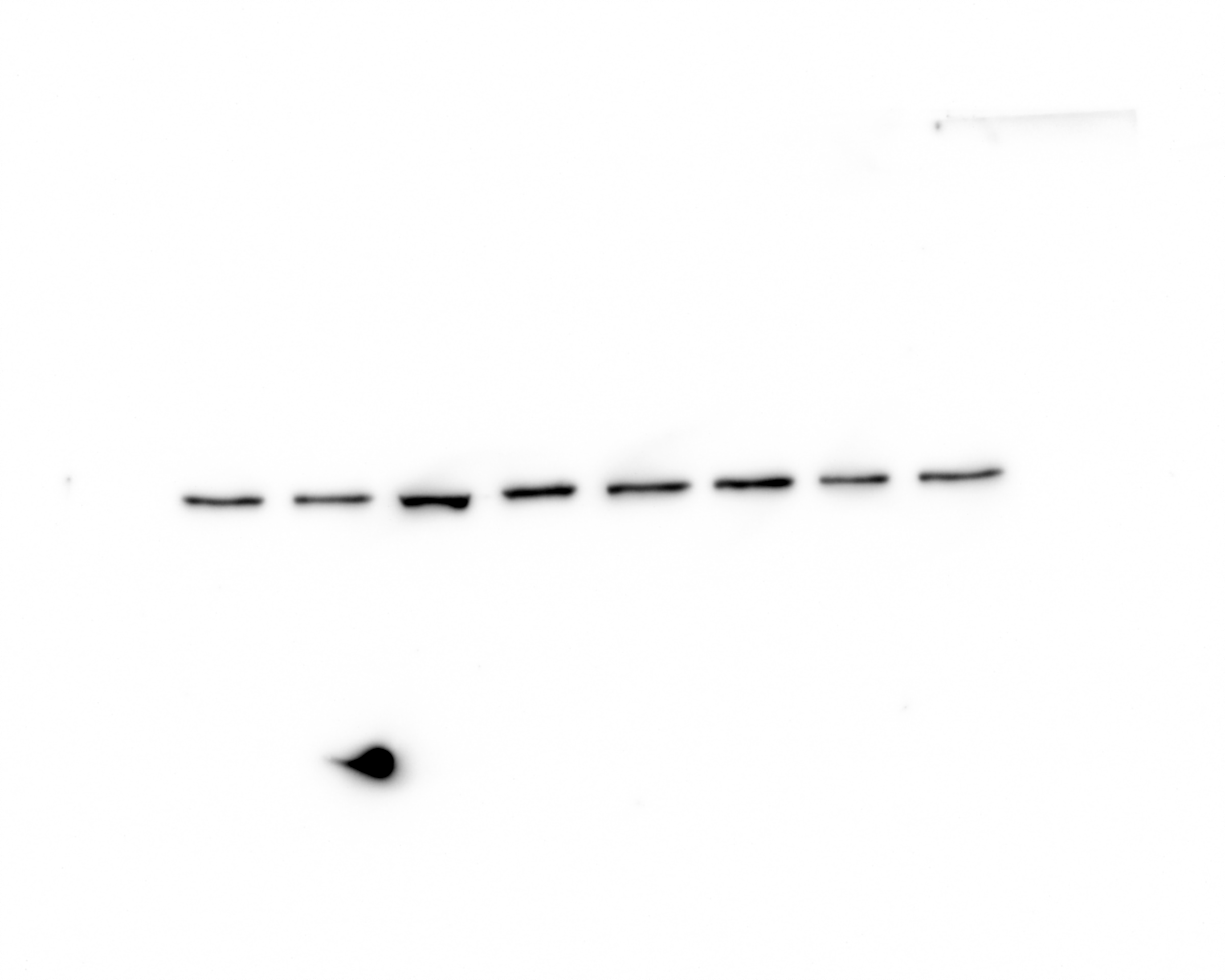

### C_msl1.tif

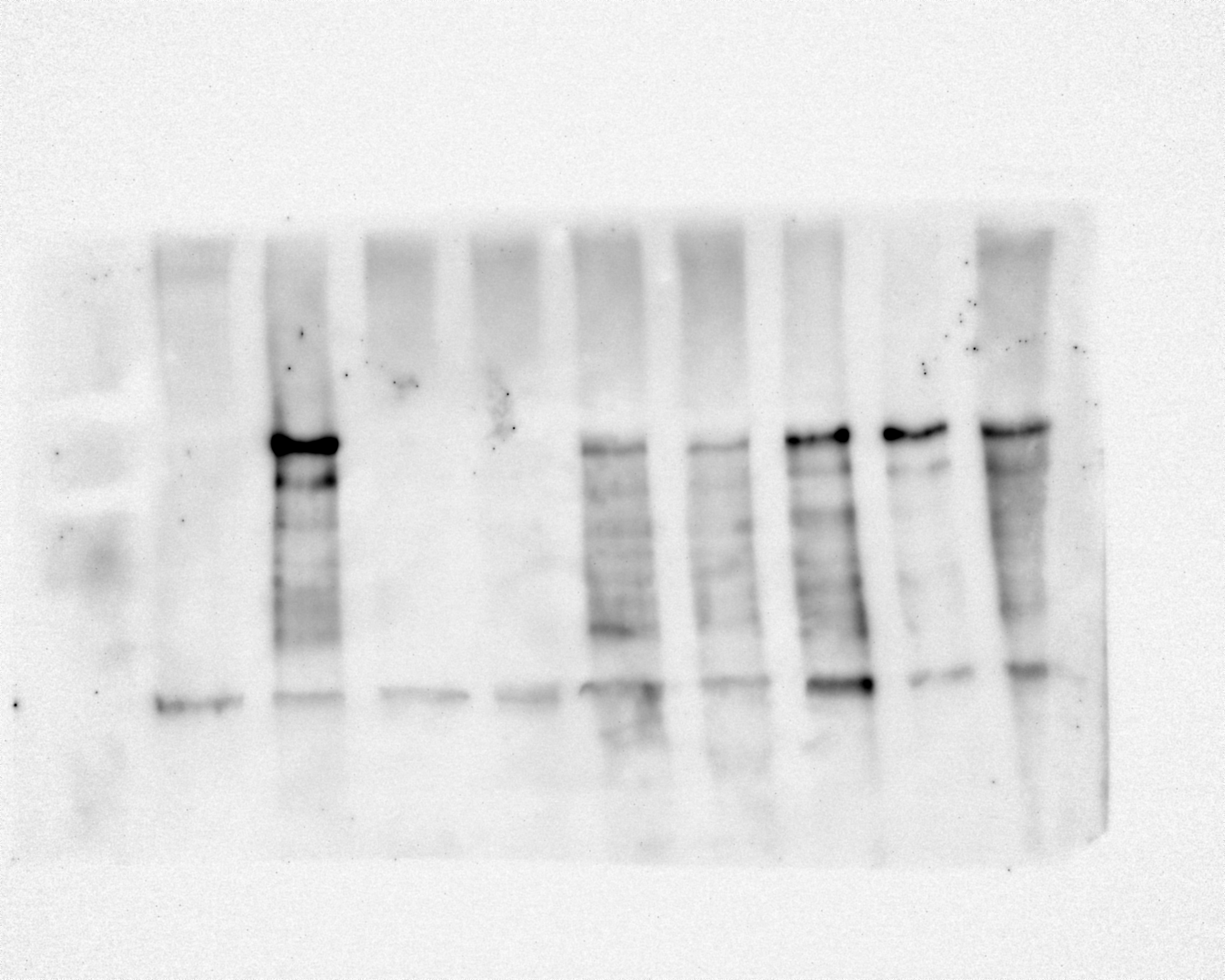

### C_msl2_m1m2_wt2a3sgs.tif

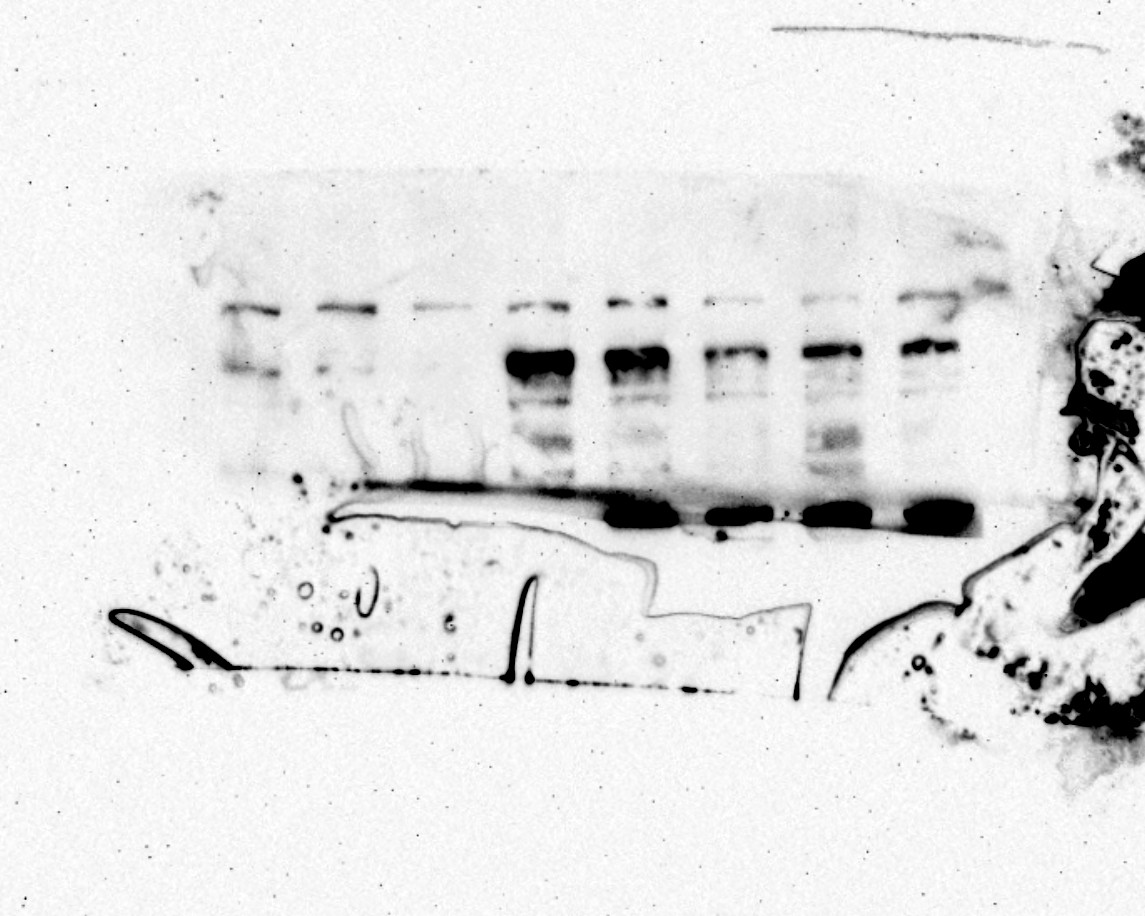

### C_msl3_m1m2_wt2a3sgs.tif

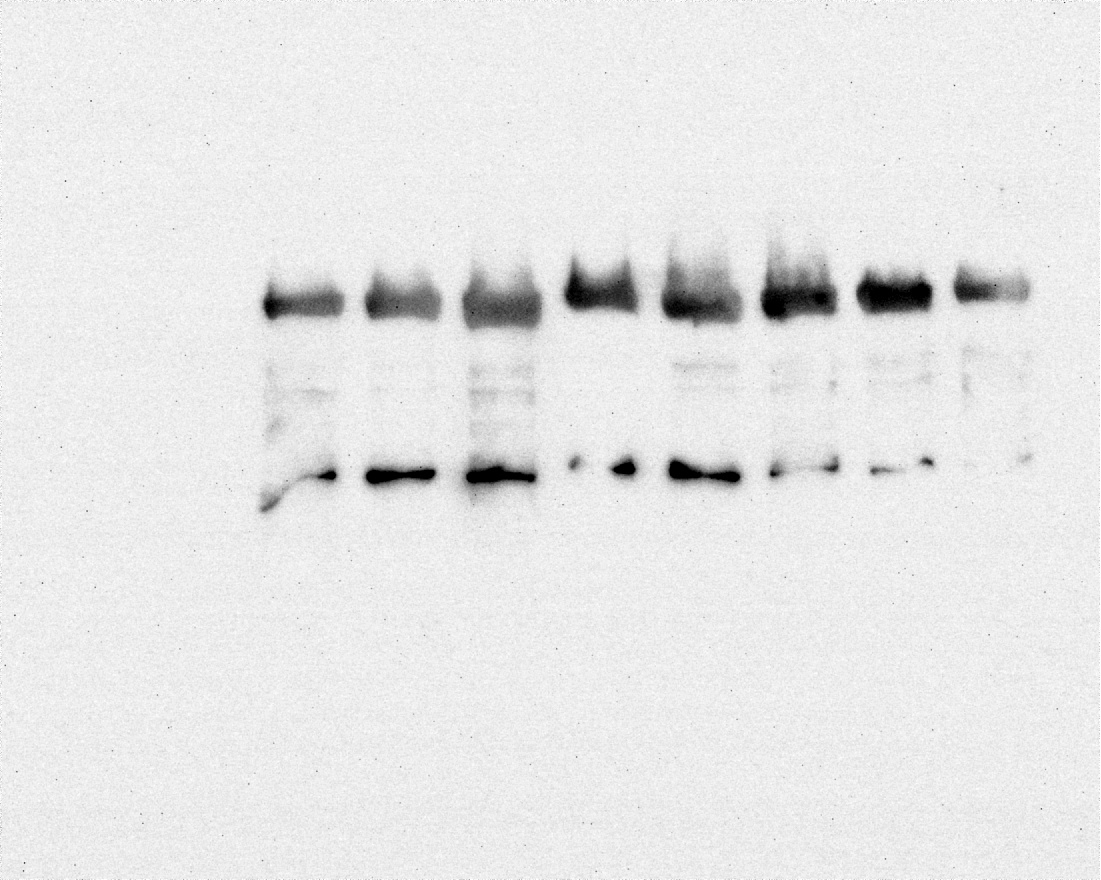

### E_lamin(Alexa 647).tif

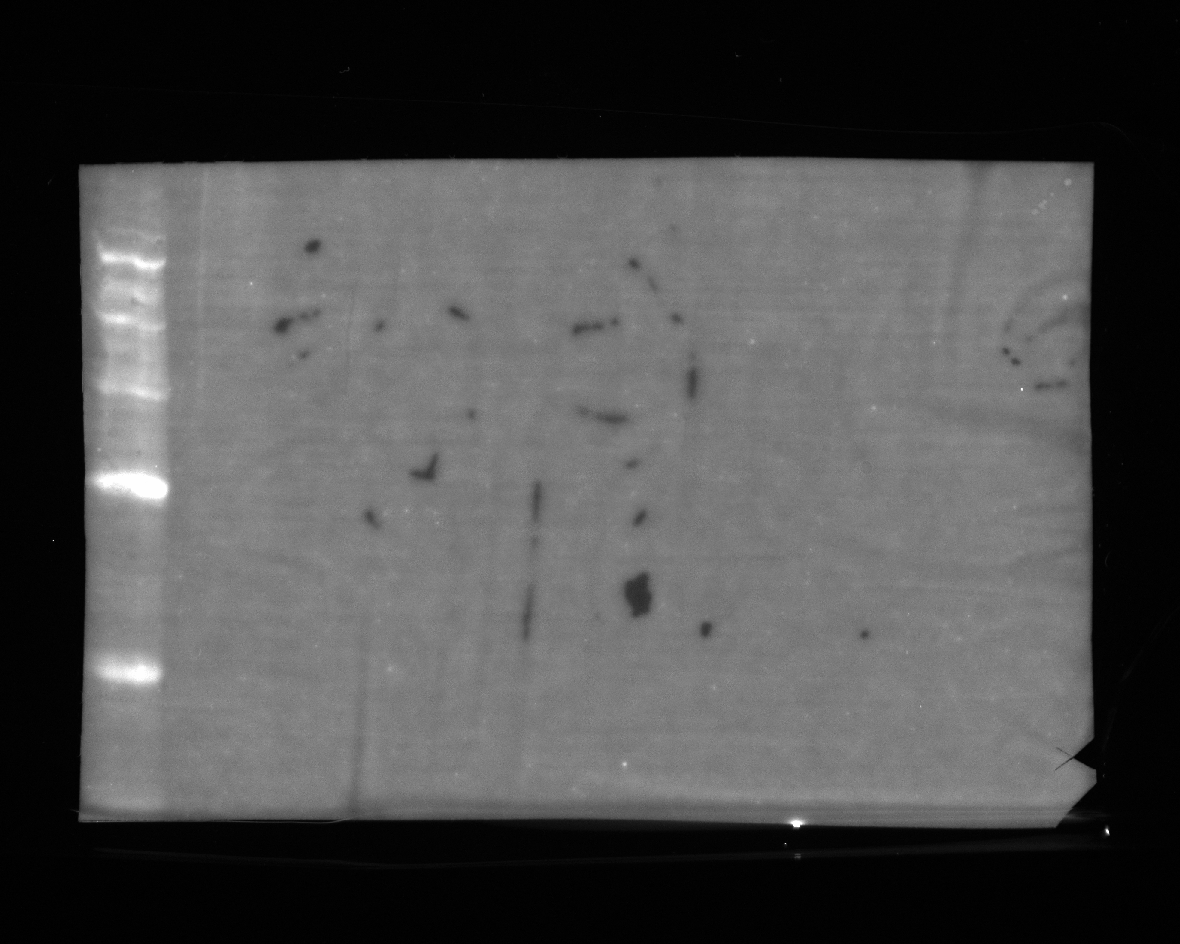

### E_lamin.tif

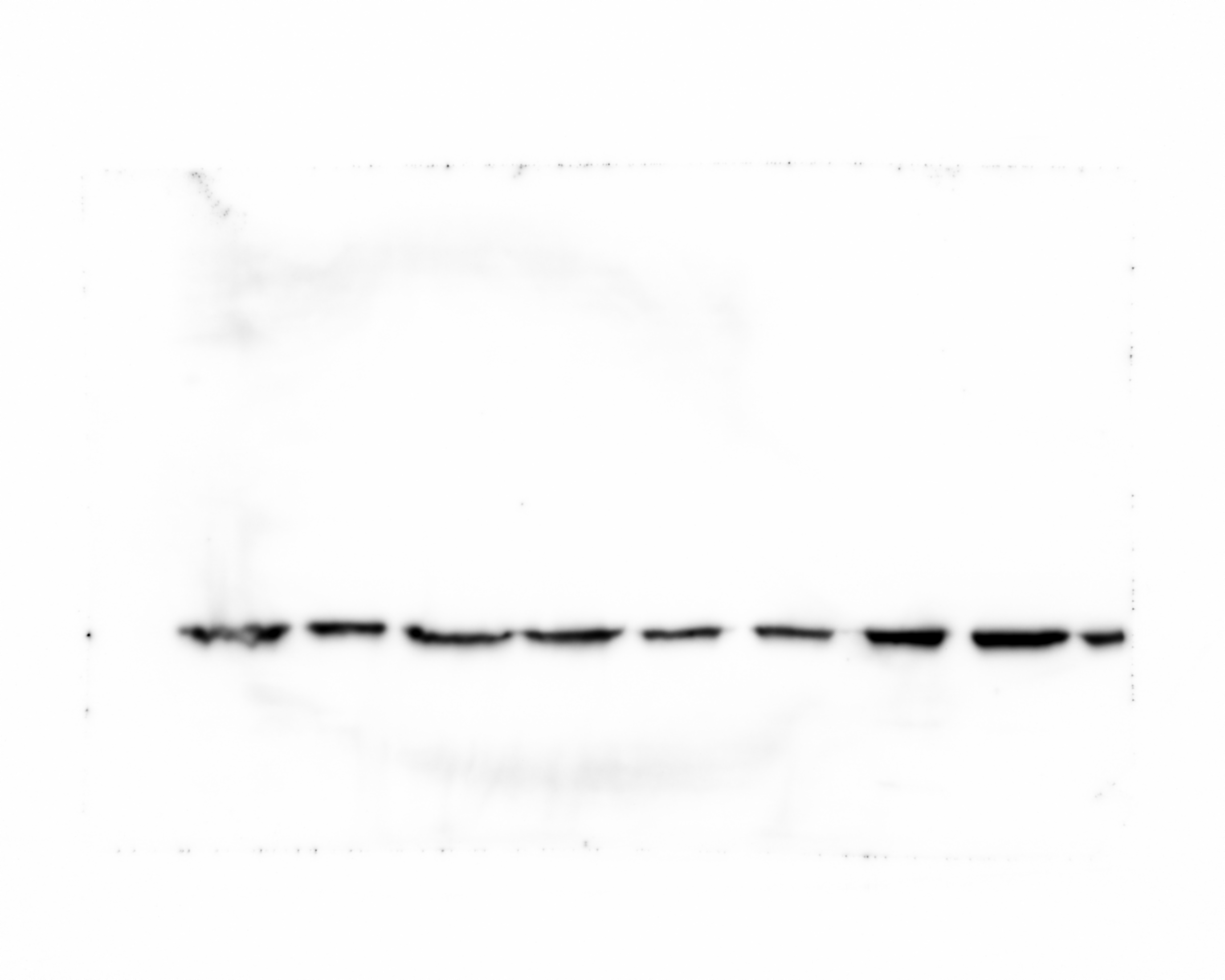

### E_msl1(Alexa 647).tif

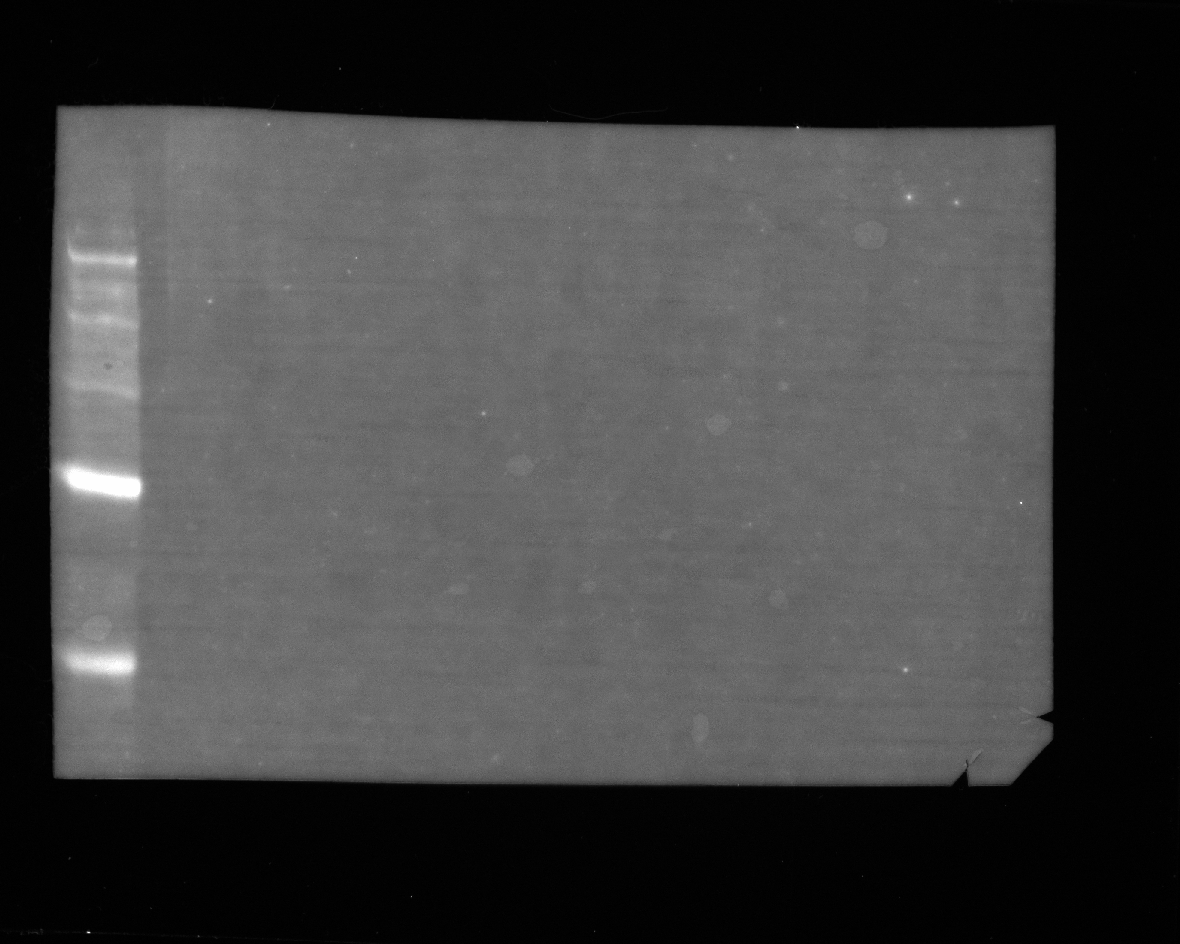

### E_msl1.tif

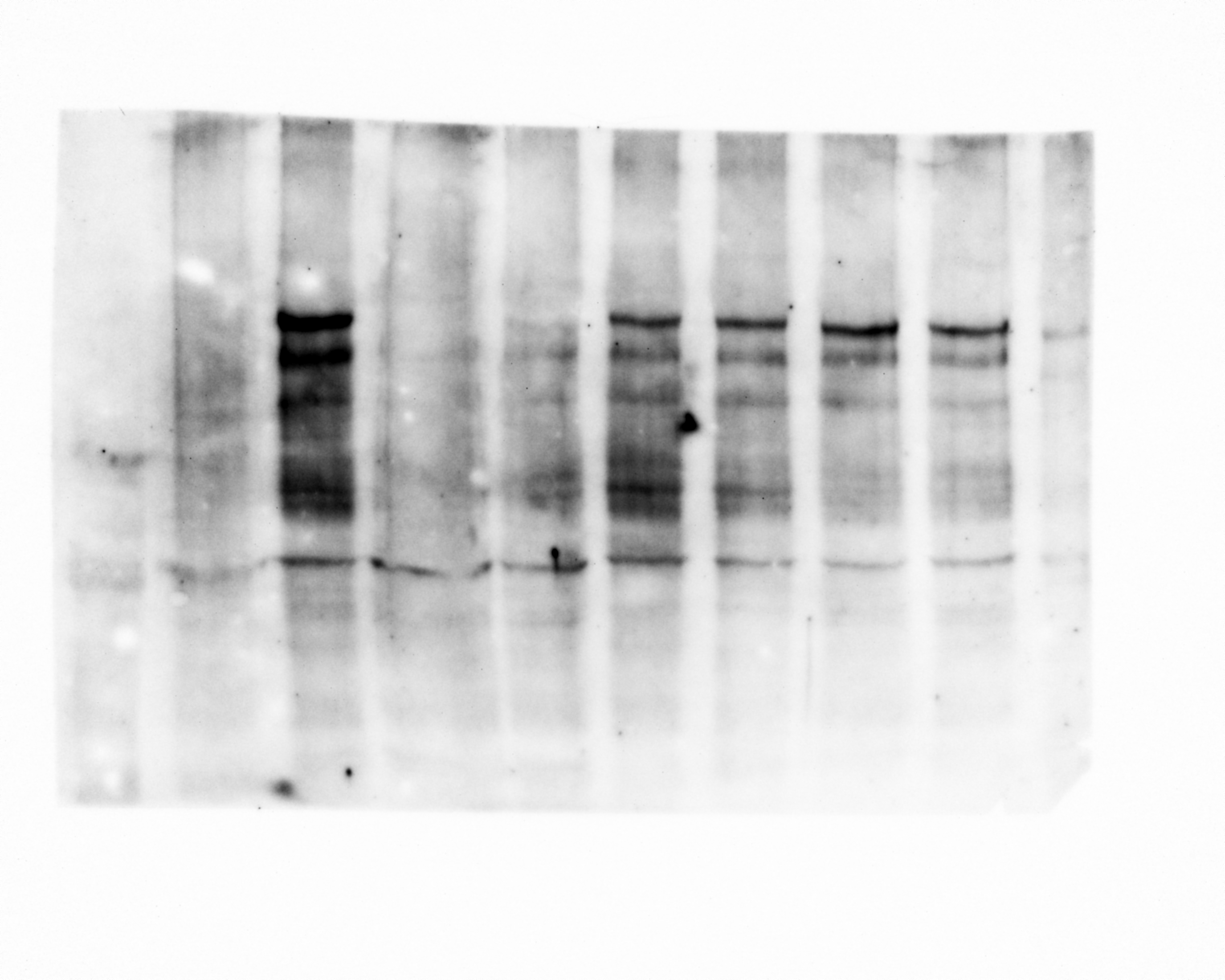

### E_msl2.tif

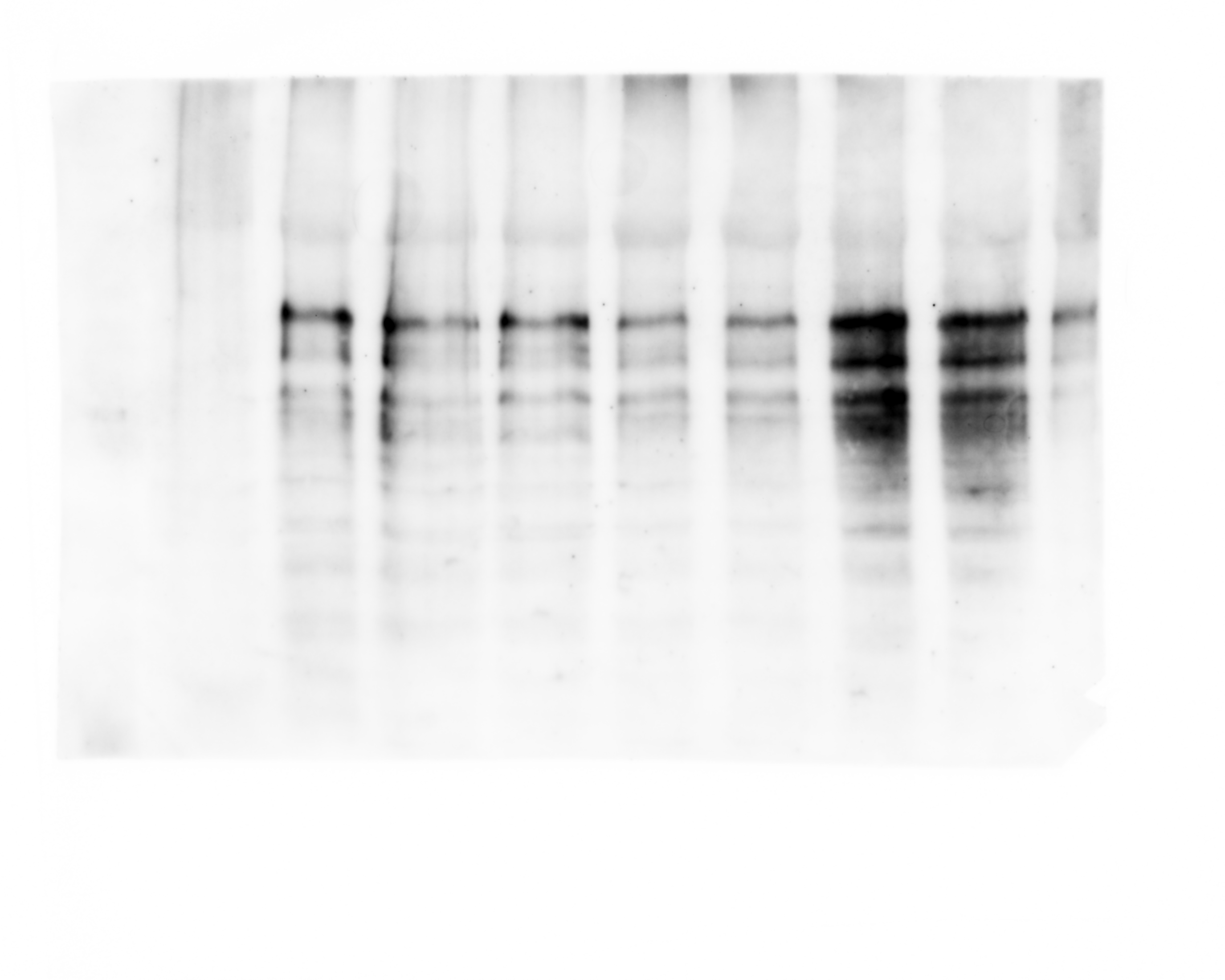
